# Trem2^hi^ macrophages bridge inflammation resolution and fibrosis initiation after ischemia-reperfusion injury in the kidney

**DOI:** 10.64898/2026.03.17.712275

**Authors:** Yan Tong, Fangxin Mu, Chao Wang, Tian Sang, Xuefeng Sun, Zhe Feng, Guangyan Cai, Xiangmei Chen, Qing Ouyang

## Abstract

Maladaptive repair of acute kidney injury (AKI) may lead to the development of chronic kidney disease (CKD) characterized by renal fibrosis. Macrophages play roles in AKI-to-CKD progression; however, the interplay between inflammation and fibrosis after AKI remains controversial and the precise role of the distinct macrophage subsets remains elusive. In the present study we identified a unique population of Trem2^hi^ macrophages derived from the bone marrow as a mediator bridging inflammation resolution and fibrosis establishment after kidney injury. Trem2 deficient mice exhibited mitigated renal fibrosis after ischemia-reperfusion injury (IRI) while the renal injury and inflammation persisted. Mechanistically, Trem2 promoted renal inflammation resolution by facilitating macrophage efferocytosis to remove apoptotic tubule cells and reshaping the macrophage cytokine production profile. Loss of Trem2 expression led to excessive cholesterol accumulation in macrophages via Lxr-Abca1/Abcg1 axis and thus sustained pro-inflammatory cytokines production. Moreover, Trem2^hi^ macrophages orchestrated the pro-fibrotic tubular epithelial cells and the activation of myofibroblasts through SPP1 to promote the establishment of renal fibrotic niche. Based on our findings, Trem2^hi^ macrophages may serve as a potential therapeutic target for AKI-to-CKD in combination with anti-inflammatory remedies.

## Introduction

Acute kidney injury (AKI) is a common clinical syndrome characterized by tubular injury and death and abrupt renal function loss ^1^, carrying high mortality, morbidity and healthcare costs, Approximately 24.6% of AKI cases progress to chronic kidney disease (CKD) with irreversible loss of renal function ^2^. Maladaptive or incomplete repair after injury promotes renal fibrosis, which is the pathological foundation of AKI-to-CKD progression ^3^. Post-injury inflammation can activate the healing program of tubular cells, meanwhile initiate fibrogenesis in the interstitium ^4^. Eventually, kidney inflammation subsides as tissue repair and extracellular matrix deposition, yet the dominant force driving inflammation resolution and fibrosis genesis is intriguing. It is critical to explore the key players in the counteraction between inflammation and fibrosis and intercept AKI-to-CKD progression.

Mononuclear phagocytes, including macrophages and dendritic cells, maintain renal immune homeostasis ^5^. After ischemia-reperfusion injury (IRI), macrophages strikingly infiltrate in the kidney and become the dominant immune cell population in inflammation response ^6,7^. The prominent recruitment of macrophages correlates with injury severity in human AKI biopsies^8^. Studies from IRI animal models have revealed that macrophages play versatile roles due to their heterogeneity during AKI-to-CKD ^9–12^. Upon acute injury, macrophages exhibited proinflammatory phenotype ^13^. Single-cell RNA sequencing (scRNA-seq) identifies S100a9^hi^ Ly6c^hi^ macrophages as a pivotal population for initiating and amplifying acute renal inflammation post-IRI ^14^, while arginase-1 expressing macrophages are recognized as a pro-reparative subset that promotes tubular proliferation^15^. However, the key macrophage populations mediating subsequent inflammation to fibrosis transition remain elusive. Thus, identifying the particular macrophage population is highly important for developing targeted therapy for AKI-to-CKD.

In this study, we identified Trem2^hi^ macrophages as an important population which bridged AKI-to-CKD transition. On one hand, Trem2^hi^ macrophages resolved renal inflammation through cholesterol-dependent manner and promoted dead tubule cells clearance through enhanced efferocytosis. On the other hand, Trem2^hi^ macrophages-derived SPP1 facilitated the establishment of fibrotic niche by activating tubular epithelial cells and myofibroblasts. These findings suggest that the pro-fibrotic and anti-inflammatory Trem2^hi^ macrophages could be a novel therapeutic target for AKI-to-CKD.

## Methods

### Animals and unilateral ischemia-reperfusion injury (uIRI) mouse model

All animal studies were conducted according to ethical guidelines approved by Animal Ethics Committee of Chinese PLA General Hospital. Male C57BL/6J mice (6-8 weeks, 20-25g body weight) were purchased from SPF Biotechnology (Beijing, China). C57BL/6J Smoc-Trem2^em1Smoc^ (Trem2KO) were purchased from Model Organisms (Shanghai, China). All the experimental animals were bred in specific-pathogen-free conditions at PLA General Hospital Animal Center.

Unilateral ischemia-reperfusion injury (uIRI) model was established to simulate AKI-to-CKD progression as previously described ^57^. After anesthetization, mice were maintained at 37 during surgery. The left renal pedicle of mouse was clamped for 35min to achieve ischemia, then the clamp was released to allow reperfusion. The contralateral kidney remained intact and functional. The right nephrectomy was performed 24h before the mouse was sacrificed to collect samples. The injured kidneys were harvested at designated time points. In the sham control group, male and age-matched mice received the identical incision and kidney exposure without renal pedicle clamping. Six mice were randomized into each group.

### Flow Cytometry Cell Sorting and scRNA Sequencing

Single cells from uIRI kidneys were stained with Zombie NIR viability dye (BioLegend) and PE-conjugated anti-mouse CD45. FSC and SSC parameters were set to exclude cell debris or mass and Zombie dye was used to exclude dead cells. For scRNA-seq, fluorescence-activated cell sorting (FACS)-isolated live CD45 and CD45□ singlets were mixed at a 4:1 ratio and resuspended at 700–1200 cells/μl with >90% viability. Singlets were captured, barcoded, and processed into libraries using the 10× Genomics Chromium system, followed by sequencing of pooled libraries on the Illumina NovaSeq6000 with 150 paired-end reads.

### Bone Marrow Transplantation Model

Recipient (CD45.2 or Trem2KO) male mice were given enrofloxacin-supplemented acidified water one week before total body irradiation (7.5Gy, Co60 source). For bone marrow (BM) transplantation, fresh BM cells were obtained from donor mice (CD45.1, WT or Trem2KO) and adoptively transferred via tail vein into irradiated recipient mice. After 4 weeks reconstruction, peripheral blood samples were collected to determine the substitution rate. And then the mice were subjected to uIRI surgery.

### Statistical analysis

Numeric data were presented as mean±SD. Unpaired Stutent’s *t* test was conducted for comparisons between two groups, and one-way analysis for variance was conducted for multiple comparisons among three or more groups. All the statistical analyses were performed using GraphPad Prism 9.1. *P* values < 0.05 were considered statistically significant.

## Results

### Immune Cell characteristics of the uIRI induced AKI-to-CKD transition

To reveal the immune landscape during AKI-to-CKD transition, we generated uIRI mice modeling the process of renal injury and fibrosis. The mice were sacrificed at 1, 3, 7, 14 days post-uIRI to represent the acute injury/inflammation stage (D1), transitional stage (D3-D7) and chronic fibrosis stage (D7-D14), respectively. Serum creatinine (Scr) and blood urea nitrogen (BUN) were evaluated to reflect renal function. The levels of Scr and BUN in uIRI mice were significantly elevated at one day post-uIRI (D1), and remained high until D7. Although slightly decreased by D14, renal function stayed subnormal due to repair failure and fibrosis genesis (Fig 1A).

**Figure 1.**
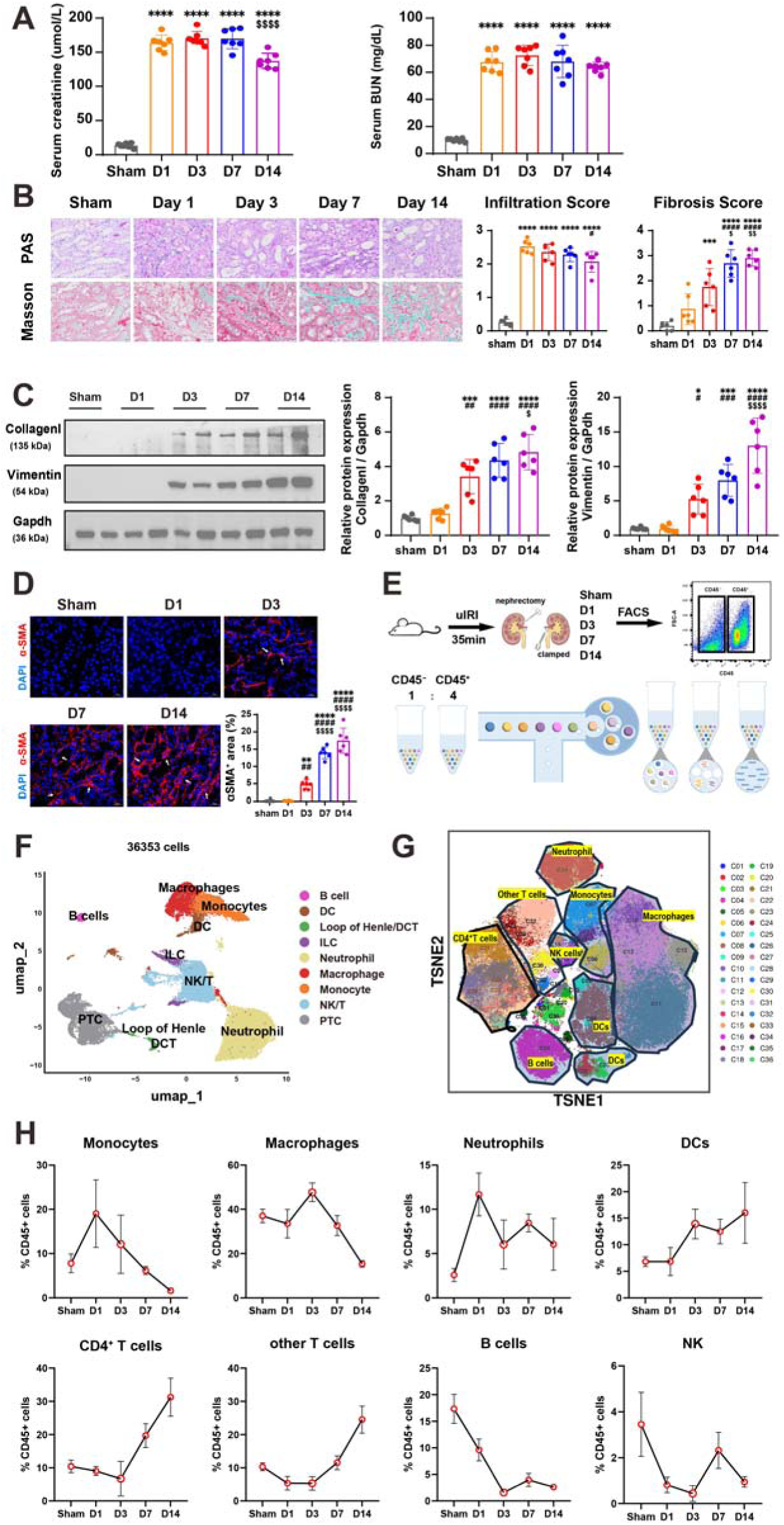
Single-cell transcriptome profiling in a murine model of AKI-to-CKD. (**A**) The levels of Scr and BUN in mice after uIRI induced AKI-to-CKD models. Representative images of PAS and Masson staining of kidney samples at different timepoints after uIRI, along with infiltration score and fibrosis score. Scale bar= 50 μm. (**B**) Expression of Collagen I and Vimentin in AKI-to-CKD mice kidney measured by western blotting. (**C**) Representative images and quantitative data of α-SMA by immunofluorescent staining. Scale bar=20 μm. (**D**) Diagram showing the experimental procedure of 10× Single-cell RNA sequencing. (**E**) Unsupervised clustering displaying kidney cell and immune cell clusters in Umap plot using single-cell RNA sequencing. (**F**) The t-SNE visualization of major immune cell subsets identified by cytometry by time of flight (CyTOF). (**G**) The percentages of each immune cell cluster in CD45^+^ cells using CyTOF across all timepoints. Data are presented as mean±SD, n=6-7, \**P* < 0.05, \*\**P* < 0.01, \*\*\**P* < 0.001, \*\*\*\**P* < 0.0001. The * represented other groups versus sham, the # represented other groups versus D1, and the $ represented other groups versus D3. Scr, serum creatinine; BUN, blood urea nitrogen; uIRI, unilateral ischemia-reperfusion injury; PAS, periodic acid-Schiff; FACS, fluorescence-activated cell sorting; PT, proximal tubule; DCT, distal convoluted tubule; DC, dendritic cell; ILC, innate lymphoid cell. CyTOF, cytometry by time of flight.

PAS staining revealed significant tubular necrosis and brush border loss at D1, with tubular regeneration and rearrangement commencing at D3. Inflammatory infiltration initiated from D1 and persisted throughout fibrotic niche formation. Masson’s Trichrome Staining indicated fibrosis onset at D3 with progressive extracellular matrix (ECM) deposition through D14 (Fig 1B). Consistently, increased expression of myofibroblast activation marker (α-SMA) and ECMs components (CollagenI, Vimentin) were detected since D3 to D14 (Fig 1C-D). These results demonstrated that inflammation response was triggered after uIRI and persisted with fibrosis formation.

Next, time series of scRNA-seq was performed on the kidneys of uIRI mice to comprehensively investigate the cellular characteristics at different stages of AKI-to-CKD transition (Fig 1E). FACS-sorted CD45 and CD45 cells from uIRI and sham kidneys were mixed at a 4:1 ratio for sequencing. After cell filtration and quality control, a total of 36,353 cells were divided into 9 clusters, including 2 clusters of kidney cells and 7 clusters of immune cells (Fig 1F). The commonly used marker genes for each cell type were illustrated in Fig S1A (Table S1). Kidney cells included primary tubules (PTs), Loop of Henle and distal convoluted tubules (DCTs). The immune cell clusters consisted of monocytes, macrophages, neutrophils, dendritic cells (DCs), B cells, NK/T cells, innate lymphoid cells (ILCs).

To validate the dynamic patterns of renal immune cell populations during AKI-to-CKD transition, we conducted mass cytometry time-of-flight (CyTOF) analysis of kidneys ^16^. Unsupervised clustering of CD45^+^ leukocytes revealed 8 distinct immune cell subsets characterized by unique cell surface marker (Fig 1G, S1B), and quantified their compositional dynamics across sequential timepoints. (Fig 1H). Under homeostasis, renal interstitium maintained macrophage residency with minimal neutrophil and monocyte infiltration. Post-injury, rapid neutrophil migration peaked at D1 then declined, likely due to their short lifespan within renal tissue. Monocytes emerged as early responders, showing significant D1 expansion concurrent with inflammatory initiation before gradual contraction. Notably, macrophages showed biphasic dynamics: initially reduced at D1, succeeded by progressive repopulation from D3 and gradual decrease by D14 when fibrosis was established, suggesting their pivotal role in orchestrating the inflammation-fibrosis axis. Dendritic cells (DCs) increased from D1 to peak at D14, despite a transient decline during D3-D7; whereas T cells declined until D3 before rising continuously through D14. B cells and natural killer (NK) cells exhibited obvious reduction since IRI without rehabitation to baseline until D14. CyTOF characterized renal immune cell population kinetics and revealed myeloid predominance during acute injury to fibrosis.

### Trem2^hi^ macrophages are a critical population mediating AKI-to-CKD transition

To identify the critical cell population in AKI-to-CKD transition, we computationally deconvolved mononuclear phagocytes (MPCs) from scRNA-seq data. Unsupervised clustering of 8,838 MPCs using Seura with Louvain optimization resolved 19 transcriptionally distinct subsets (Fig 2A, S2A, table S2), systematically annotated through marker gene expression profiling ^14,17^. Resident macrophage populations (Clusters 2/3/7/9) were identified by conserved markers (C1qa/b/c, Cd81, Ms4a7) with subset-specific signatures: Cluster 2 (Egr1^hi^), Cluster 3 (MHCII^hi^), Cluster 7 (Rpl^hi^), and Cluster 9 (Mrc1^hi^). Infiltrating macrophages (Clusters 4/5/10/17) exhibited dichotomous polarization aligning with M1/M2 paradigms (Fig 2B) ^18^. Pro-inflammatory subsets comprised CCR2^hi^ (Cluster 5) and S100A8/9^hi^ (Cluster 17) populations, while reparative subsets included Arg1^hi^Fn1^hi^ (Cluster 4) and Trem2^hi^Spp1^hi^ (Cluster 10) macrophages. Temporal analysis (Fig 2C) revealed distinct macrophage dynamics across AKI-to-CKD progression. Resident macrophages (Clusters 2, 3, 7, 9) were profoundly depleted upon initial injury (D1), indicating susceptibility to initial damage. Although some exhibited transient recovery, most failed to fully reconstitute, indicating a permanent loss of the homeostatic pool. On the other hand, infiltrating populations (Clusters 4, 5, 10, 17) progressively expanded during acute inflammation phase and remained prominent in the established fibrotic phase, highlighting their important role in mediating the transition.

**Figure 2.**
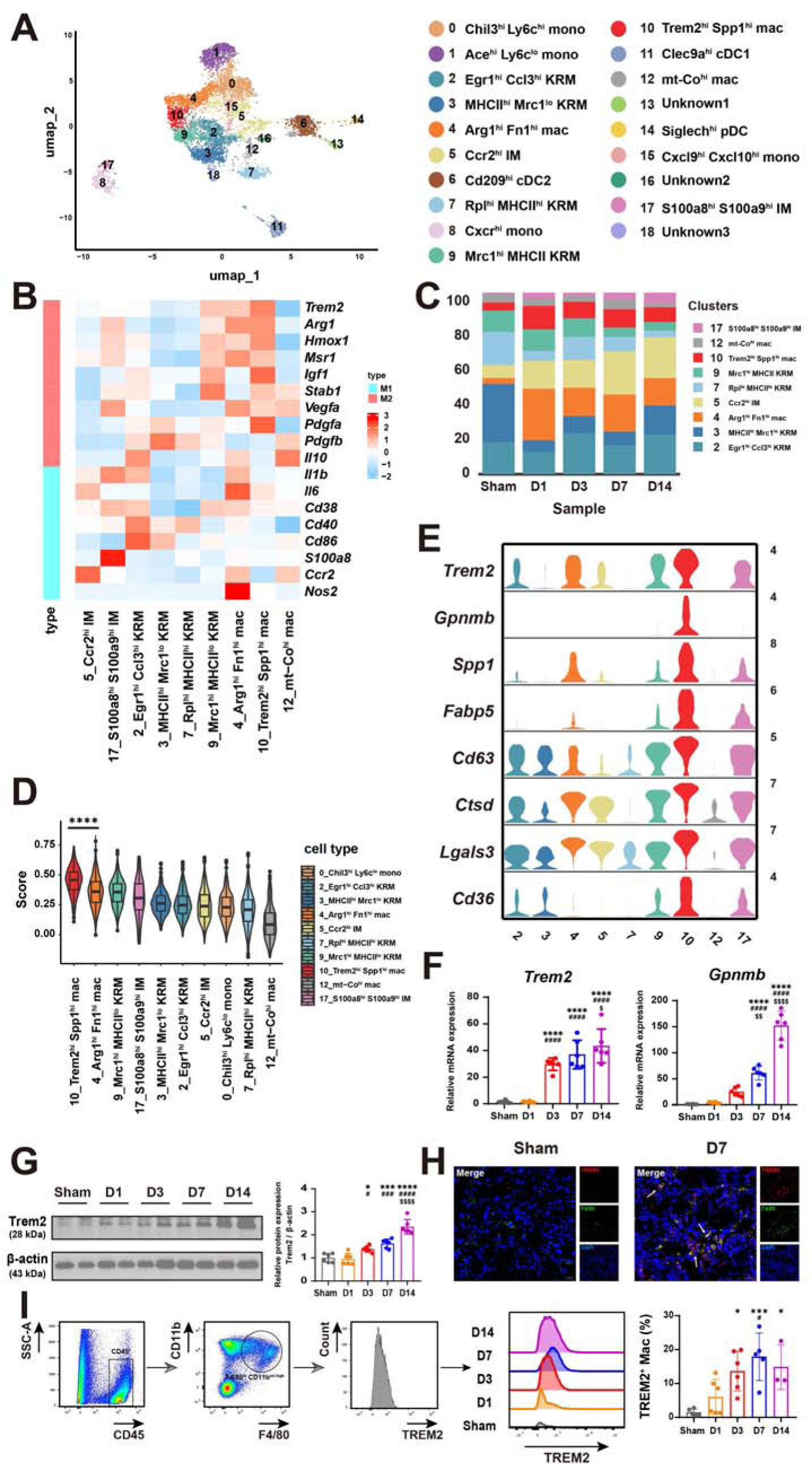
scRNA-seq analysis identified discrete MPC clusters and important Trem2^hi^ macrophages. **(A)** Sub-clustering of MPC clusters of all samples and MPC annotation. **(B)** Heatmap of the expression of classical M1 or M2 macrophage marker genes in all macrophage clusters across all samples. **(C)** Bar graphs showing the relative proportions of respective macrophage clusters. **(D)** ECM regulator scores of macrophage clusters across all samples. **(E)** Volin plot exhibiting the expression of Trem2 and associated genes in respective macrophage cluster across all samples. **(F)** Relative mRNA expression level of Trem2 and Gpnmb in AKI-to-CKD mice kidney measured by RT-qPCR. **(G)** Expression of Trem2 protein level in AKI-to-CKD mice kidneys measured by western blotting. **(H)** Immunofluorescence images of Trem2^+^ macrophages in sham and D7 kidney. **(I)** Percentage of Trem2^hi^ macrophages in AKI-to-CKD kidney detected by flow cytometry. Data are presented as the mean±SD, n=6, \**P* < 0.05, \*\**P* < 0.01, \*\*\**P* < 0.001, \*\*\*\**P* < 0.0001. The * represented other groups versus sham, the # represented other groups versus D1, and the $ represented other groups versus D3.

Notably, Trem2^hi^ macrophages emerged as a fibrogenesis-associated subpopulation co-expressing reparative and profibrotic transcripts. Functional analysis using Matrisome Project ^19^-defined ECM regulatory gene sets demonstrated superior fibrotic potential in Trem2^hi^ macrophages versus other subsets (Fig 2D), positioning them as pivotal regulators of AKI-to-CKD progression. We therefore identified its defining marker Trem2 as a key gene. Expression of Trem2 was also observed in clusters 4 and 9 (Fig S2B), which correspondingly displayed elevated fibrotic scores (Fig 2F). This consistent correlation led us to hypothesize that Trem2 expression in macrophages is associated with their fibrotic potential. To confirm the significance of Trem2^hi^ macrophages in AKI-to-CKD progression, we performed multimodal validation. Core signature genes (*Trem2, Lgals3, Spp1, Gpnmb, Cd63, Ctsd, Cd36, Fabp5*) showed progressive transcriptional activation of *Trem2* and *Gpnmb* initiating at D3 by RT-qPCR (Fig 2E-F). Western blotting demonstrated sustained Trem2 protein upregulation from D3 through D14, mirroring fibrotic progression kinetics (Fig 2G). Spatial characterization via immunofluorescence confirmed the interstitial localization of F4/80^+^Trem2^hi^ macrophages (Fig 2H). Flow cytometry analysis of the F4/80^hi^CD11b^int-high^ compartment revealed that the proportion of Trem2^+^ macrophages was minimal in Sham groups but progressively increased following uIRI during D3-D7, and slightly decreased by D14 (Fig 2I, S2C). Collectively, these results established Trem2^hi^ macrophages as critical effector coordinating the inflammation-fibrosis axis in AKI-to-CKD transition.

### Trem2 knockout ameliorates fibrosis induced by uIRI

To elucidate whether Trem2^hi^ macrophages could affect AKI-to-CKD progression, we generated Trem2 knockout (KO) mice (Fig S3A-B) and compared their responses with WT mice following uIRI. Initially, KO mice exhibited attenuated functional impairment, with lower Scr at D7 and reduced BUN at D7 and D14 versus WT mice (Fig 3A). Histological evaluation via Masson’s Staining revealed comparable baseline fibrosis of sham groups both in WT and KO mice, however, KO mice exhibited significantly lower fibrosis scores at D7 and D14 (Fig 3B). Consistently, KO mice showed significantly decreased staining of fibrotic marker CollagenI than WT mice (Fig 3C). Western blotting analysis further confirmed decreased protein levels of CollagenI and Vimentin in KO kidneys (Fig 3D), while RT-qPCR showed downregulation of *Col1a1*, *Fn1* and *Tgfb* (Fig S3C). These data suggested that Trem2 deficiency attenuates renal fibrosis at D7 and D14 after injury

**Figure 3.**
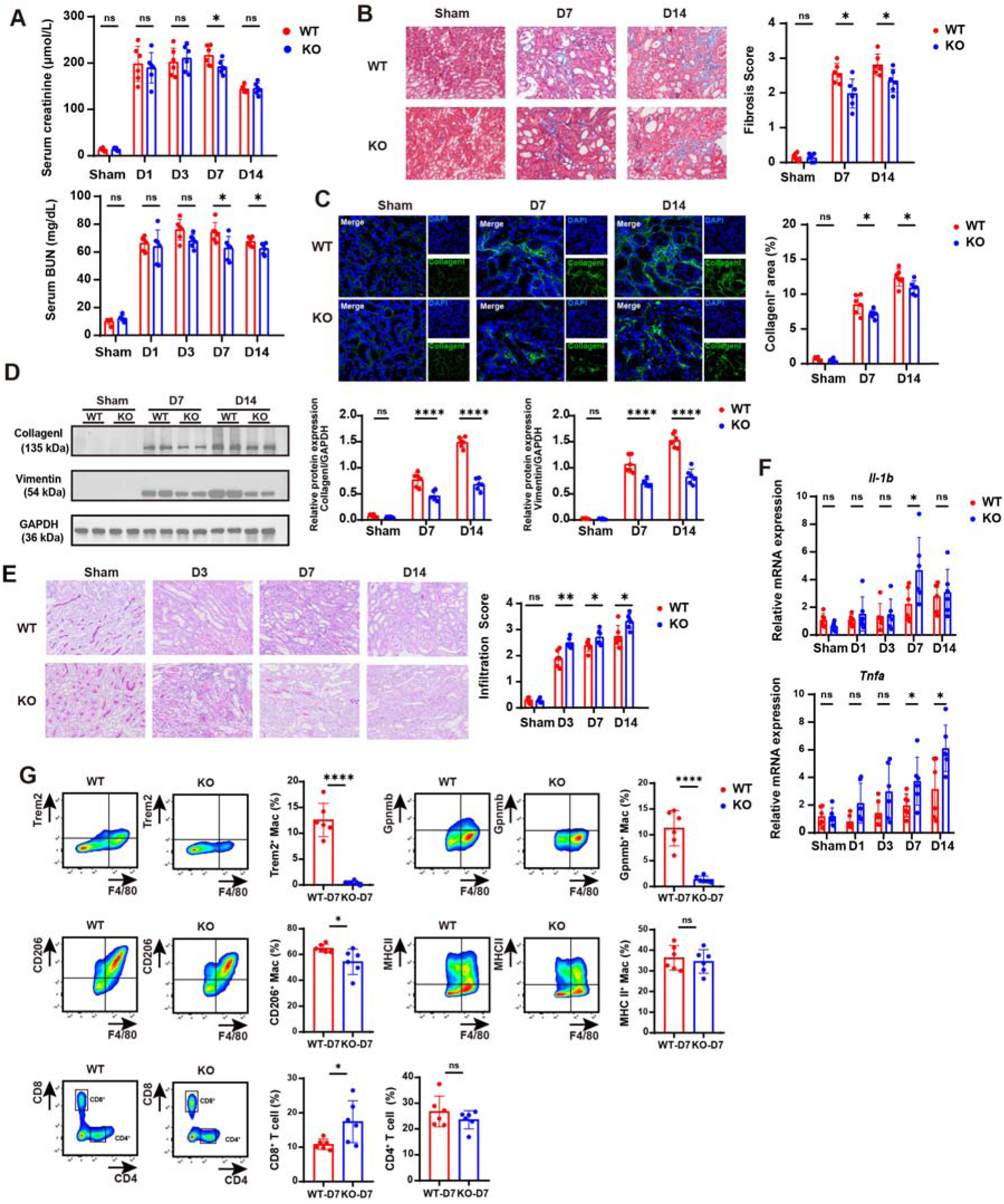
Trem2 knockout alleviated fibrosis, but aggravated inflammation after uIRI. **(A)** The levels of Scr and BUN in WT and KO mice of AKI-to-CKD models. **(B)** Masson staining of kidney samples WT and KO mice. Scale bar= 50 μm. **(C)** Representative images and quantitative data of Collagen I by immunofluorescent staining. Scale bar= 20 μm. **(D)** Expression of Collagen I and Vimentin in WT and KO mice kidneys measured by western blotting. **(E)** PAS staining of kidney samples WT and KO mice. Scale bar= 50 μm. **(F)** Relative mRNA expression of inflammatory cytokines detected by RT-qPCR. **(G)** The percentages of other immune cells in WT and KO mice at D7 were analyzed by flow cytometry. Data are presented as the mean±SD, n=6, \**P* < 0.05, \*\**P* < 0.01, \*\*\**P* < 0.001, \*\*\*\**P* < 0.0001, ns, no significance. PAS, periodic acid-Schiff; Fn1, fibronectin; *Il-1b*, interleukin 1β; *Tnfa*, tumor necrosis factor; mac, macrophage.

We then assessed the effect of Trem2 deficiency on inflammation. In terms of inflammation scores, there were no differences between WT and KO mice at homeostasis. However, at D3, D7 and D14 after injury, higher inflammation scores were observed in KO mice than WT (Fig 3B). Although inflammatory cytokines (*Il-1b*, *Tnfa*) mRNA increased in both groups after uIRI, KO mice manifested prolonged expression peaks (Fig 3F), aligning with the sustained inflammation in PAS staining. These results suggested that Trem2 deletion aggravated non-resolved inflammation at least until D14 after uIRI.

To interrogate how Trem2 deficiency modified the immune microenvironment during their regulatory phase in AKI, we analyzed immune cell populations in the kidneys of uIRI mice at D7 using flow cytometry. Trem2^hi^ macrophages constituted 12.58% of F4/80^hi^ CD11b^int-high^ macrophages in kidneys of WT mice, the number in KO mice is 0.46%. Considering that Gpnmb was one of the top feature genes for Trem2^hi^ macrophages, we quantified Gpnmb^+^ macrophages and found them similarly significantly reduced in KO mice. The proportion of anti-inflammatory CD206^+^ macrophages also decreased from 64.83% in WT to 54.37% in KO mice, indicating mitigated inflammation resolution in KO kidneys. The percentages of MHCII^+^ macrophages showed no difference between WT and KO mice. The reduction of Trem2^hi^ macrophages and their associated anti-inflammation/pro-regenerative macrophage populations (Gpnmb^+^ and CD206^+^) may contribute to sustained injury and inflammation in KO mice. Lymphoid population analysis revealed comparable percentages of CD19^+^ B cells and CD4^+^ T cells between WT and KO mice. Notably, KO mice exhibited a significant increase of renal CD8^+^ T cells, consistent with the findings in tumor microenvironment ^20^, indicating that Trem2 deficiency not only impaired myeloid-mediated inflammation restriction, but also enhanced adaptive immune-driven injury (Fig 3G, Fig S3E-F). The absolute numbers of specific subsets per 200,000 total cells were presented in Fig S3G.

### Trem2^hi^ macrophages are derived from bone marrow

As shown in the scRNA-seq data (Fig S2), Trem2^hi^ macrophages expressed some resident macrophages markers (C1qa, C1qb and Cd81). To determine their origin, yolk-sac-derived resident macrophages proliferation or infiltrated monocytes differentiation, we performed RNA velocity analysis across all timepoints (Fig 4A). Results showed that Ly6c^hi^ monocytes (Cluster 0) differentiated into Arg1^hi^Trem2^lo^ macrophages (Cluster4) at D1 post-injury, which subsequently evolved into Trem2^hi^Spp1^hi^ macrophages (Cluster 10) with upregulated Trem2 expression.

**Figure 4.**
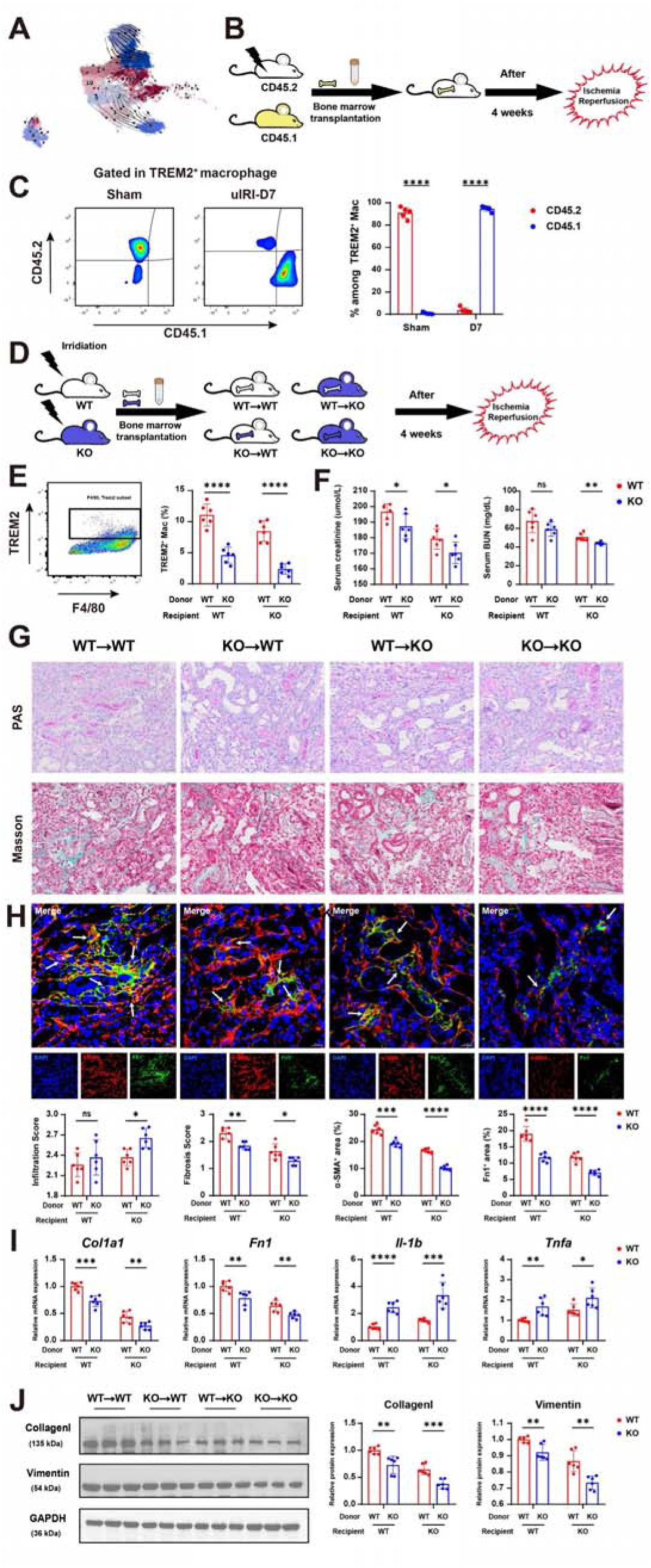
The origin and function of Trem2^hi^ macrophages during AKI-to-CKD. **(A)** Umap plot of kidney MPC clusters showed the developmental transition as revealed by RNA velocity. **(B)** Experimental protocol of BMT to elucidate the origin of Trem2^hi^ macrophages. **(C)** CD45.1 or CD45.2 expression of Trem2^hi^ macrophages after uIRI at D7 were analyzed by flow cytometry. **(D)** Experimental protocol of BMT to confirm the function of Trem2^hi^ macrophages. **(E)** Percentages of Trem2^hi^ macrophages in BMT mice were analyzed by flow cytometry. **(F)** The levels of Scr and BUN in BMT mice. **(G)** Representative images of PAS and Masson staining of kidney samples in BMT mice, along with inflammation score and fibrosis score. Scale bar= 50 μm. **(H)** Representative images and quantitative data of α-SMA, and Fn1 by immunofluorescent staining. Scale bar= 20 μm. **(I)** Relative mRNA expression levels of *Col1a1*, *Fn1*, *Il-1b* and *Tnfa* were measured by RT-qPCR. **(J)** Expression of Collagen I and Vimentin in BMT mice measured by western blotting. Data are presented as the mean±SD, n=5-6, \**P* < 0.05, \*\**P* < 0.01, \*\*\**P* < 0.001, \*\*\*\**P* < 0.0001, ns, no significance. PAS, periodic acid-Schiff; *Col1a1*, Collagen I; *Fn1*, fibronectin; *Tgfb*, transforming growth factor β; *Il-1b*, interleukin 1β; *Tnfa*, tumor necrosis factor.

To track the origin of Trem2^hi^ macrophages after uIRI, we established bone marrow transplantation (BMT) models. Briefly, Trem2^+/+^ CD45.2 mice were irradiated to remove their bone marrow (BM) cells, and received BM cells of Trem2^+/+^ CD45.1 mice, in which the macrophages from circulation were CD45.1^+^ and resident cells in the kidneys were CD45.2^+^. After 4 weeks reconstitution, the BMT mice were subjected to uIRI and the composition of CD45.1^+^ and CD45.2^+^ compartments within Trem2^hi^ macrophages in the kidneys were analyzed by flow cytometry at D7 (Fig 4B, Fig S4B). In the healthy kidneys, 91% of Trem2^hi^ macrophages were CD45.2^+^, and less than 1% were CD45.1^+^, indicating that the Trem2^hi^ macrophages were resident macrophages at homeostasis. However, at D7 post-uIRI, nearly 95% of Trem2^hi^ macrophages were CD45.1^+^, and CD45.2^+^ Trem2^hi^ macrophages reduced to 3% (Fig 4C). Our data confirmed the bone marrow origin of fibrotic Trem2^hi^ macrophages, indicating a dramatic replacement of the resident population. The vulnerability of resident macrophages to acute injury enables repopulation by newly recruited macrophages, which subsequently proliferate and become the long-term contributors to the fibrotic niche.

To validate the direct role of bone marrow-derived Trem2^hi^ macrophages in AKI-to-CKD transition, we generated another BMT model (Fig 4D). Four chimeric groups were generated by transplanting WT or KO bone marrow into irradiated WT and KO recipients (donor→recipient): WT→WT, WT→KO, KO→WT, KO→KO. After 4 weeks, the chimeric mice underwent uIRI. Based on our characterization of the dynamic progression of AKI-to-CKD, we focused on D7 post-uIRI as a pivotal phase where established inflammation coincides with evident fibrosis to compare macrophage populations across BMT groups. Flow cytometry revealed that recipients of WT BM (WT→WT, WT→KO) showed significantly more Trem2^hi^ macrophages than those receiving KO BM (KO→WT, KO→KO) (Fig 4E, S4C). The highest proportion occurred in WT→WT mice, while KO→WT mice exhibited significant reduction, indicating the Trem2^hi^ macrophages were mostly cleared in KO→WT mice. The more Trem2^hi^ macrophages in WT→KO mice compared with those in KO→KO mice confirmed their successful migration from bone marrow to injured kidney post-uIRI.

Transplantation of KO BM showed improved renal function (Fig 4F) and attenuated fibrosis (Fig 4G) compared with their counterparts receiving WT BM. We also noticed an elevation of inflammation score in the kidneys of KO→KO mice compared with those in the WT→KO mice, although the comparison bewttwen KO→WT and WT→WT mice was not significant (Fig 4G). In parallel with the results of Masson staining, the expression of fibrotic markers (α-SMA, TGF-β, Collagen I, Fn1 and Vimentin) increased in WT→WT, WT→KO mice compared with those in KO→WT, KO→KO mice (Fig 4H-J, Fig S4E). The replenishment of Trem2^hi^ macrophages restored the mitigated fibrosis in KO mice, which supported the pro-fibrotic function of Trem2^hi^ macrophages.

Mice receiving KO BM (KO→WT and KO→KO) showed negligible Trem2 mRNA (Fig S4D) and elevated inflammatory cytokines (*Il-1b*, *Tnfa*, *Ccl2*) versus WT BM recipients (Fig 4I, Fig S4E), suggesting Trem2 deficiency promoted a pro-inflammatory state of macrophages. KO BM recipients showed specifically reduced CD206^+^ macrophages but increased CD8^+^ and T-bet^+^ T cells, without affecting B cells, Foxp3^+^ or PD-1^+^ T cells (Fig S4F-G). These results suggested that recruitment of Trem2-deficient macrophages into the kidney could mitigate fibrosis and prolong inflammation to a later stage, indicating that Trem2^hi^ macrophages bridging the transition from inflammation to fibrosis.

### Trem2 deficiency impairs efferocytotic clearance of dead tubule cells by macrophages and aggravates subsequent renal inflammation

To decipher the specific functions of Trem2^hi^ macrophages in AKI-to-CKD, we identified 47 significantly upregulated genes in Trem2^hi^ macrophages based on the scRNA-seq data (Fig 5A). KEGG pathway analysis showed that these genes were enriched in lysosome, phagosome and cholesterol metabolism (Fig 5B), while GO analysis highlighted enrichment in collagen-containing ECMs and ECMs binding (Fig S5A). These transcriptomic signatures suggested multifaced roles for Trem2^hi^ macrophages in cellular debris clearance, metabolic reprogramming and fibrotic niche formation.

**Figure 5.**
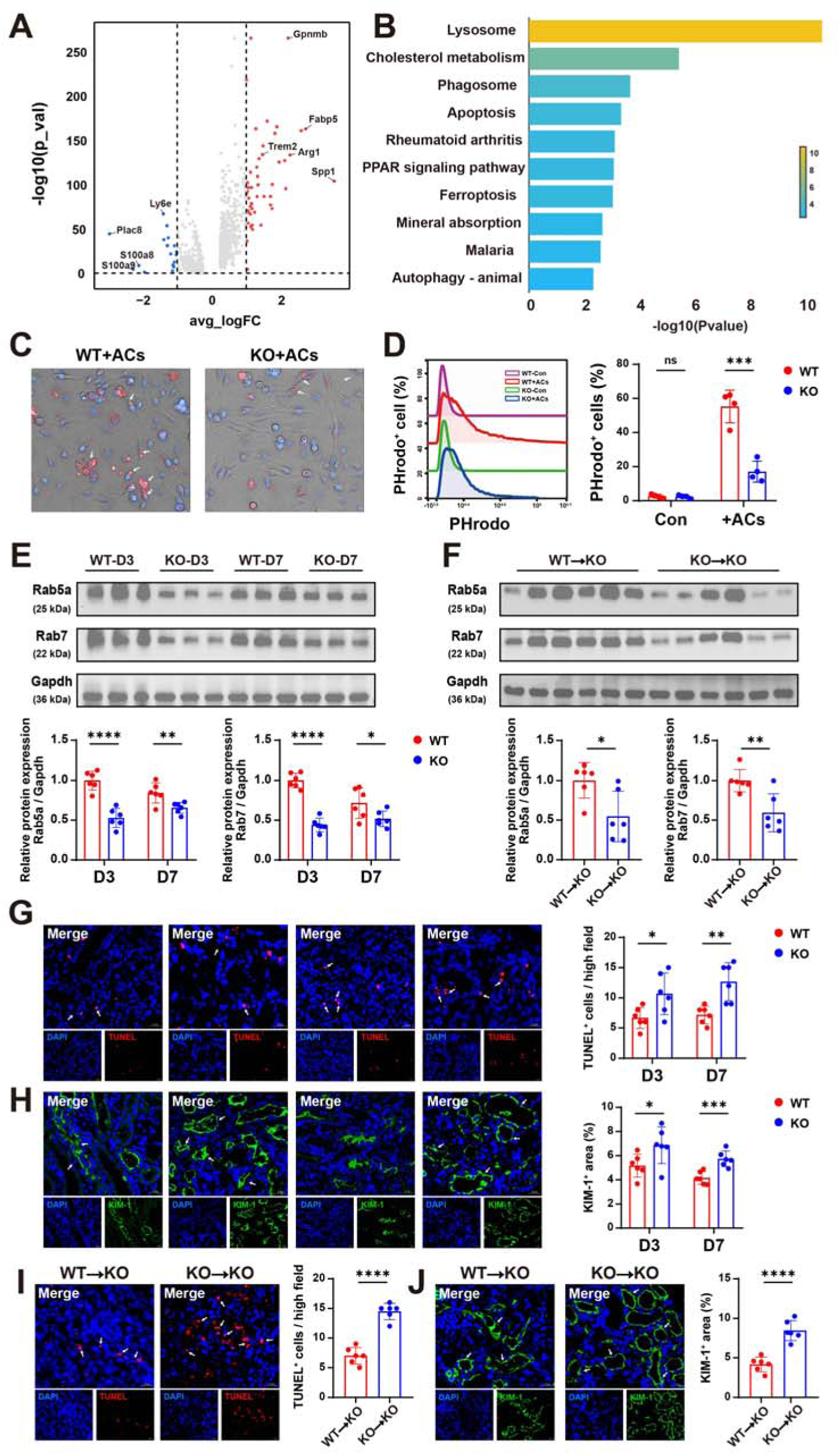
Trem2 is essential for macrophage efferocytosis to remove apoptotic cells. **(A)** Volcano plot displaying the differentially expressed genes between Trem2^hi^ macrophages and other MPCs in uIRI kidneys. **(B)** Top 10 enriched KEGG terms of Trem2^hi^ macrophages in injured kidneys. **(C)** Representative images of WT and KO BMDMs to uptake PHrodo+ ACs. **(D)** The efferocytosis rates of WT and KO BMDMs to uptake PHrodo+ ACs detected by flow cytometry. **(E)** Western blotting of Ras-related protein Rab5a and Rab7 in WT and KO mice kidneys at D3 and D7 post-uIRI. **(F)** Western blotting of Rab5a and Rab7 in BMT model of WT→KO and KO→KO mice kidneys at D7 post-uIRI. **(G)** Immunofluorescence staining images of uIRI kidney sections from WT and KO mice with TUNEL (red) and DAPI (blue). Scale bar=20 μm. **(H)** Representative images and quantitative data of KIM-1 (green) from WT and KO kidneys by immunofluorescent staining. Scale bar= 20 μm. **(I)** Immunofluorescence staining images of uIRI kidney sections from WT→KO and KO→KO mice with TUNEL (red) and DAPI (blue). Scale bar=20 μm. **(J)** Representative images and quantitative data of KIM-1 (green) from WT→KO and KO→KO kidneys by immunofluorescent staining. Scale bar= 20 μm. Data are presented as the mean±SD, for in vitro experiments, n=4, for in vivo experimetns, n=6. \**P* < 0.05, \*\**P* < 0.01, \*\*\**P* < 0.001, \*\*\*\**P* < 0.0001, ns, no significance. Rab5a, Rab7, Ras-related proteins; KIM1, kidney injury molecule 1; Con, control; ACs, apoptotic cells.

As shown in previous studies, I/R-induced AKI triggers tubular cell apoptosis ^21^. Efficient clearance of apoptotic cells (efferocytosis) by macrophages is critical for inflammation resolution and tissue repair ^22,23^. To assess the efferocytotic capacity of Trem2^hi^ macrophages indicated in transcriptome, we co-cultured bone marrow-derived macrophages (BMDMs) from WT or KO mice with apoptotic tubule cells in vitro. Tubular cell hypoxic injury was modeled using rotenone, a mitochondrial complex inhibitor, which induced apoptosis in 53.7 % of cells (Fig S5B). Apoptotic cells (ACs) were labeled with PHrodo Red, a pH-sensitive fluorescent dye that fluoresces red in acidic environments upon lysosomal internalization. Flow cytometry revealed that Trem2 deficiency severely impaired efferocytotic activity, with only 17% of KO-BMDMs being PHrodo^+^ versus 55% of WT (Fig 5C-D). Lysosome-associated membrane protein (LAMP) expression quantitatively reflects efferocytosis efficiency by marking functional lysosomal compartmentalization required for phagocytic flux ^23^. Immunofluorescence showed WT-BMDMs exhibited significantly higher basal LAMP expression than KO-BMDMs, and both increased after ACs stimulation. However, WT-BMDMs remained superior LAMP expression (Fig S5C). The mRNA expression level of Trem2 was negligible in KO-BMDMs (Fig S5D). This pronounced impairment demonstrates that Trem2 signaling is indispensable for efferocytosis, providing mechanistic explanation for the unresolved inflammation and delayed tissue repair in Trem2-deficient mice during AKI-to-CKD.

We next examined phagocytic activity in vivo through evaluating the expression of endosomal markers in injured kidneys. Trem2-WT mice exhibited significantly higher expression of early (Rab5a) and late (Rab7) endosomal markers compared to Trem2-KO mice (Fig 5E). BMT experiments also confirmed the phagocytosis of Trem2^hi^ macrophages, as Rab5a/Rab7 levels were restored in WT→KO versus KO→KO chimeras (Fig 5F), demonstrating bone marrow-derived Trem2^hi^ macrophages drive the process of phagocytosis in injured kidneys. To clarify the consequences of failed efferocytosis, we performed TUNEL staining to indicate the ACs in kidneys. Trem2-KO kidneys displayed more TUNEL^+^ ACs than Trem2-WT kidneys (Fig 5G), confirming defective clearance. While WT BM transplantation rescued this impairment (Fig 5I), directly linking Trem2^hi^ macrophages to ACs removal. Concurrently, Trem2-KO mice exhibited prolonged renal injury evidenced by more KIM-1+ tubular cells (Fig 5H, 5J) and elevated KIM-1 mRNA expression (Fig S5E) at D3/D7 post-injury. In BMT experiments, KO→KO chimeras showed greater tubule injury than WT→KO counterparts (Fig 5J, S5E). These findings demonstrated that effective Trem2^hi^ macrophages-mediated efferocytosis is essential for prevention of sustained renal injury.

### Trem2 deficiency causes cholesterol accumulation and enhances inflammatory response in macrophages

As indicated by KEGG (Fig 5B), Trem2^hi^ macrophages is associated with cholesterol metabolism. Given that intracellular cholesterol overload can lead to inflammasome activation and IL-1β-driven inflammation ^24^, we investigated the role and underlying mechanisms of Trem2 signaling in cholesterol metabolism and inflammation. BODIPY (Boron-Dipyrromethene) staining revealed higher baseline lipid deposition in KO-BMDMs than WT-BMDMs. After exposure to ACs, both groups showed increased lipid deposition, yet KO-BMDMs maintained significantly higher deposition than WT-BMDMs (Fig 6A). Cholesterol-targeted metabolomics revealed that KO-BMDMs had markedly elevated oxysterols (2.56-fold) despite minimal change in total sterols (Fig 6B). Consistently, Filipin staining confirmed pronounced intracellular cholesterol accumulation in KO-BMDMs (Fig 6C) compared with WT-BMDMs, indicating Trem deficiency disrupts cholesterol homeostasis.

**Figure 6.**
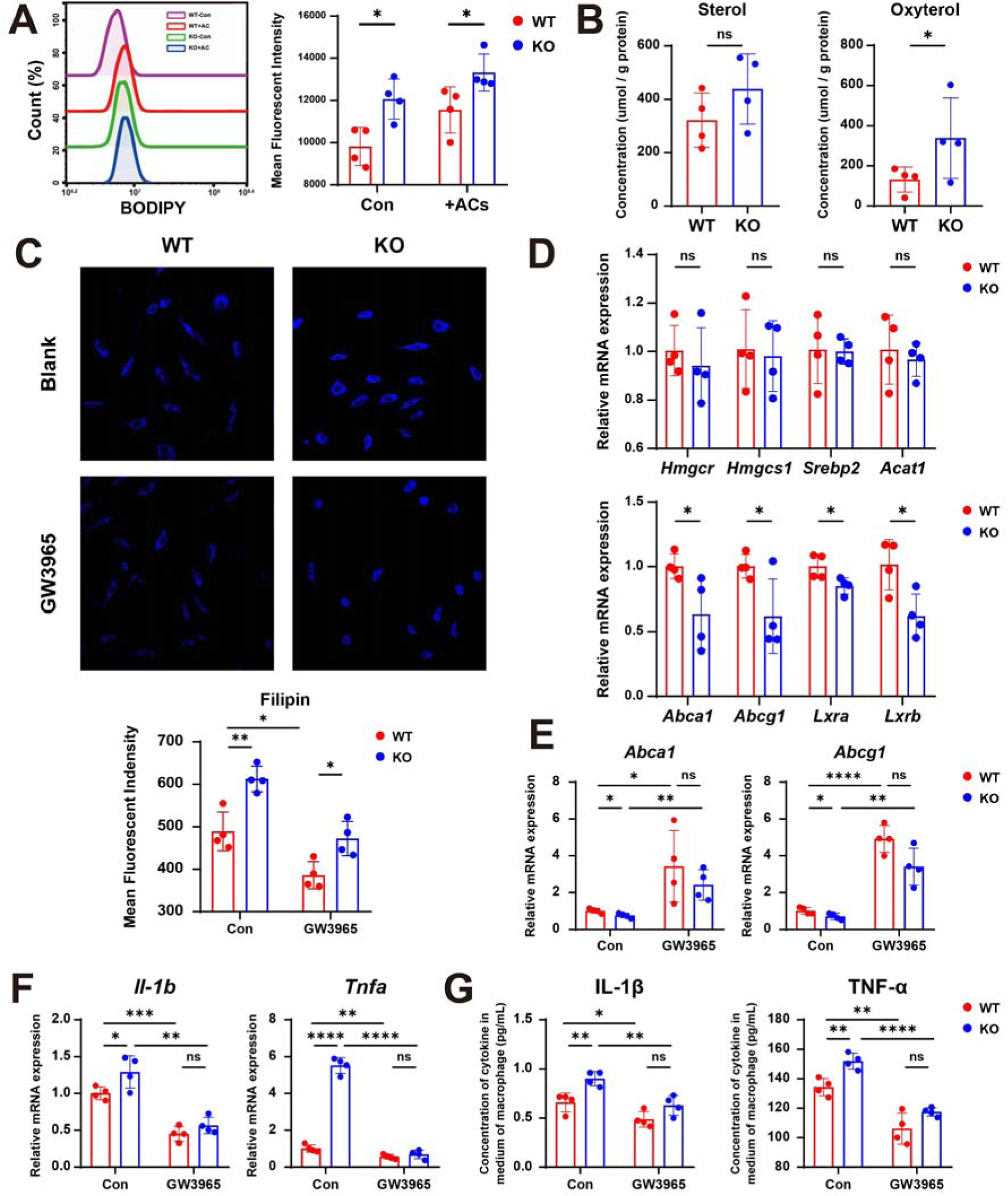
Trem2 regulated cholesterol metabolism and inflammation. **(A)** BMDMs from WT and KO mice in the absence or presence of ACs were stained with BODIPY and detected by flow cytometry. **(B)** Quantification of sterol and oxysterol between WT-BMDMs+ACs and KO-BMDMs+ACs. **(C)** The filipin staining of BMDMs from WT and KO mice in the absence or presence of LXR activation (GW3965). **(D)** Relative mRNA expression levels of cholesterol metabolism related genes were measured by RT-qPCR. **(E)** Relative mRNA expression levels of *Abca1* and *Abcg1* were measured by RT-qPCR after LXR activation (GW3965). **(F)** Relative mRNA expression levels of *Il-1b* and *Tnfa* were measured by RT-qPCR after LXR activation (GW3965). **(G)** The concentration of cytokine IL-1β and TNF-α in BMDMs from WT and KO mice in the media were measured by ELISA after LXR activation (GW3965). Data are presented as the mean±SD, n=4, \**P* < 0.05, \*\**P* < 0.01, \*\*\**P* < 0.001, \*\*\*\**P* < 0.0001, ns, no significance. Con, control; ACs, apoptotic cells; LAMP, lysosome-associated membrane proteins; *Hmgcr*, 3-hydroxy-3-methylglutaryl-Coenzyme A reductase; *Hmgcs1*, 3-hydroxy-3-methylglutaryl-Coenzyme A synthase 1; *Srebp2*, sterol regulatory element binding factor 2; *Acat1*, acetyl-Coenzyme A acetyltransferase 1; *Abca1*, ATP-binding cassette, sub-family A member 1; *Abcg1*, ATP binding cassette subfamily G member 1; *Lxra*, Liver X receptor α; *Lxrb*, Liver X receptor β.

To elucidate how TREM2 regulates cholesterol metabolism in macrophages, we detected key cholesterol metabolism regulatory genes in BMDMs. While cholesterol synthesis genes (*Hmgcr*, *Hmgcs1* and *Srebp2*) and cholesterol esterification gene (*Acat1*) were unaffected, KO-BMDMs showed marked downregulation of liver X receptors (LXRs) isoforms (*Lxra*, *Lxrb*), and their target cholesterol efflux genes (*Abca1*, *Abcg1*) which promote reverse cholesterol transport in macrophages ^25^ (Fig 6D). This transcriptional profile suggests Trem2 signaling sustains cholesterol efflux through LXR-dependent regulation of *Abca1/Abcg1*. Pharmacological activation of LXRs with GW3965 rescued the mitigated *Abca1/Abcg1* expression in KO-BMDMs, while enhancing their expression in both WT- and KO-BMDMs (Fig 6E). Consistently, GW3965 treatment reduced intracellular cholesterol accumulation in both WT- and KO-BMDMs as shown in Filipin staining and narrowed their discrepancy (Fig 6C), indicating that GW3965 partially restored the cholesterol efflux in KO-BMDMs via activating LXR-ABCA1/ABCG1 pathway.

Myeloid LXR deficiency promotes inflammatory gene expression in atherosclerotic foamy macrophages ^26^, indicating its regulatory roles in lipid metabolism and inflammation suppression ^27^. Therefore, we determined the levels of proinflammatory cytokines in WT- and KO-BMDMs. At baseline, KO-BMDMs showed elevated expression of *Il1b* and *Tnfa* than WT-BMDMs (Fig 6F). GW3965 treatment significantly suppressed these pro-inflammatory cytokines in both genotypes, particularly diminishing the differences between WT- and KO-BMDMs (Fig 6F). ELISA assay confirmed enhanced IL-1β and TNF-α secretion in KO-BMDMs compared with WT-BMDMs, which was attenuated upon LXR activation (Fig 6G). These findings collectively demonstrated that impaired LXR expression functionally contributed to excessive cholesterol accumulation and pro-inflammatory cytokine release in Trem2-deficient macrophages.

### Trem2^hi^ macrophages promote renal fibrosis by interacting with TECs and myofibroblasts via Spp1

Tubular epithelial cells (TECs) and myofibroblasts constitute key effector cells in renal fibrosis ^28^. To determine how Trem2^hi^ macrophages promote fibrosis, we performed CellChat analysis on our scRNA-seq data and integrated external datasets^29^ to survey intercellular communication networks. Four TECs subpopulations with specific gene profiles were identified: normal TECs (*Gatm, Miox*), injured TECs (*Havcr1, Fgg*), pro-fibrotic TECs (*Malat1, Neat1*) and progenitors (*Cryab, Tmsb10*) (Fig 7A, S6) ^30^. AUCell pathway scoring revealed that pro-fibrotic TECs exhibited marked enrichment in fibro-inflammatory pathways, including hypoxia, IL-6-JAK-STAT3, TNF-α signaling, inflammatory response and TGF-β signaling (Figure 7B). Intercellular communication analysis demonstrated dominant interactions between Trem2^hi^ macrophages and pro-fibrotic TEC cluster across both extracellular matrix-receptor (ER) and paracrine/autocrine signaling (SS) networks (Fig 7C). Ligand-receptor profiling identified *Spp1* (secreted phosphoprotein 1) as the predominant mediator produced by Trem2^hi^ macrophages engaging integrins (*Itgav/Itgb1/6*) on pro-fibrotic TECs (Fig 7D-E, S7A). Similar SPP1-integrin axis-driven crosstalk was observed between Trem2^hi^/Arg^hi^ macrophages and myofibroblasts (Fig 7F-G, S7B) in external scRNA-seq datasets ^29^. Immunofluorescence revealed F4/80^+^ Trem2^hi^ macrophages adjacent to α-SMA^+^ stromal cells (Fig S7C), suggesting the direct crosstalk between Trem2^hi^ macrophages and pro-fibrotic cells. These data indicated SPP1 as a master regulator of Trem2^hi^ macrophage-stromal interactions promoting renal fibrosis.

**Figure 7.**
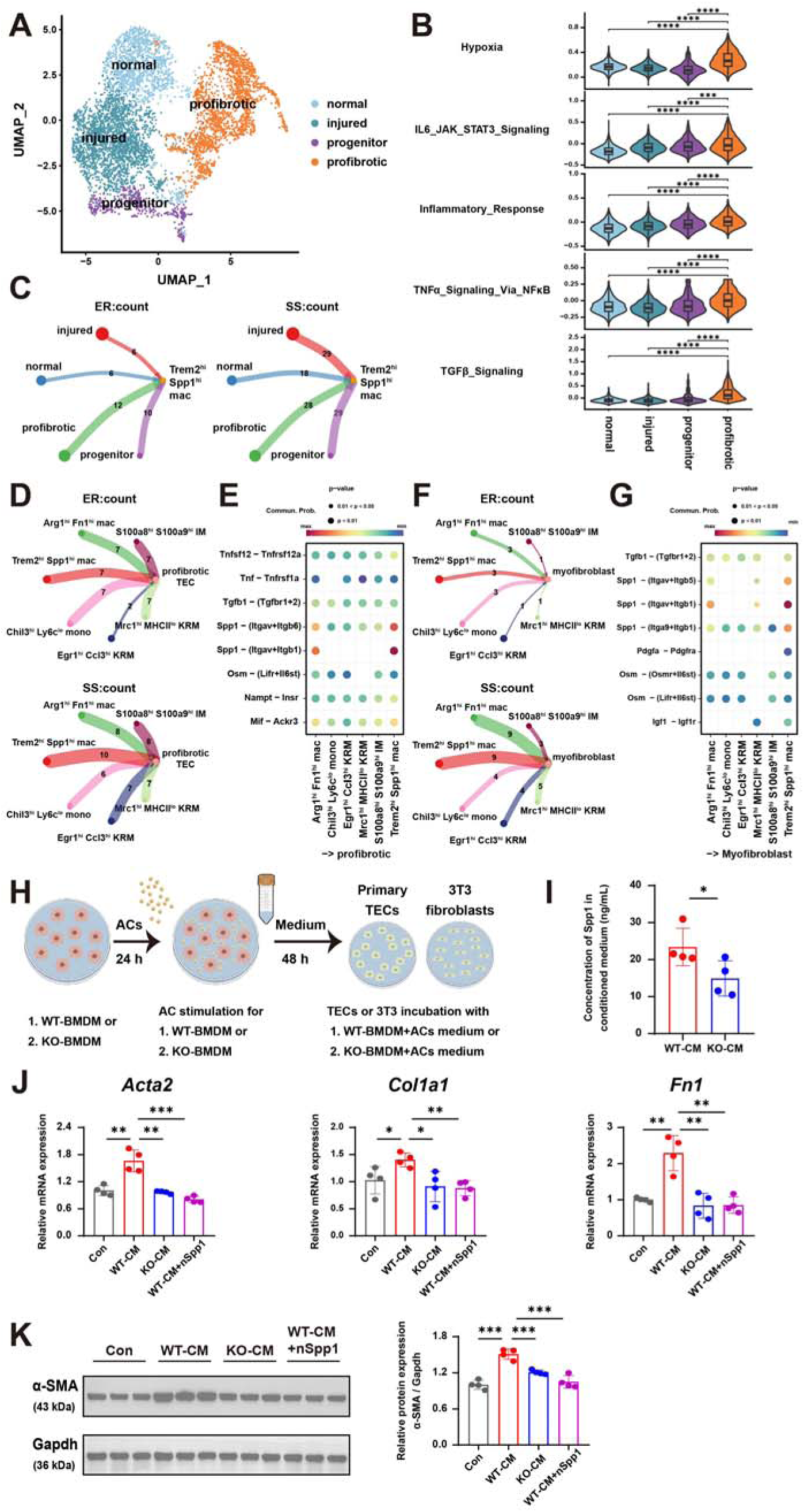
Trem2^hi^ macrophages activated TECs and fibroblasts via Spp1 to promote renal fibrosis. **(A)** Unsupervised clustering displaying kidney tubular cell clusters in Umap plot. **(B)** Violin plots showing pathway enrichment in the TEC subclusters. **(C)** The circle plots showing the numbers of interactions between TECs subtypes and Trem2^hi^ macrophages. **(D)** The circle plots showing the numbers of interactions between macrophage subtypes and pro-fibrotic TECs. **(E)** The heatmap showing the ligand-receptor interactions between macrophage subtypes and pro-fibrotic TECs. **(F)** The circle plots showing the numbers of interactions between macrophage subtypes and myofibroblasts. **(G)** The heatmap showing the ligand-receptor interactions between macrophage subtypes and myofibroblasts. **(H)** Scheme by Figdraw for the experiment design of kidney TECs and fibroblasts stimulated by the media of WT-BMDMs and KO-BMDMs incubated with ACs. **(I)** The concentration of Spp1 in CM was detected by ELISA. **(J)** RT-qPCR analysis of *Acta2*, *Col1a1* and Fn1 mRNA in primary TECs after stimulation by CM or CM+Spp1 neutralization antibody for 48 h. **(K)** Western blotting of α-SMA in 3T3 fibroblasts after stimulation by CM or CM+Spp1 neutralization antibody for 48 h. Data are presented as the mean±SD, n=4, \**P* < 0.05, \*\**P* < 0.01, \*\*\**P* < 0.001, \*\*\*\**P* < 0.0001, ns, no significance. TECs, tubular epithelial cells; mac, macrophage; KRM, kidney resident macrophage; IM, infiltrating macrophage; mono, monocyte; CM, conditioned media.

To functionally validate SPP1-mediated Trem2^hi^ macrophages-stromal cells interactions, we treated TECs and fibroblasts with conditioned media (CM) from ACs-phagocytosing WT/KO BMDMs (Fig 7H). Higher SPP1 concentrations in WT-CM than KO-CM were confirmed by ELISA (Fig 7I). The isolated primary TECs exposed to WT-CM exhibited upregulated fibrogenic genes (Acta2, Col1a1, Fn1) versus KO-CM (Fig 7J). WT-CM stimulated stronger myofibroblast activation marker α-SMA expression in 3T3 fibroblasts than KO-CM (Fig 7K). To confirm the causal role of Spp1 in the induction of profibrotic phenotype, we used Spp1 neutralization antibody in the culture system. Compared with WT-CM, the addition of Spp1-blocking antibody significantly suppressed the upregulation of fibrogenic markers in TECs by RT-qPCR (Fig 7J) and attenuated α-SMA expression in fibroblasts by western blotting (Fig 7K), demonstrating that Spp1 is dispensable for Trem2^hi^ macrophage mediated fibrosis. Collectively, based on these results, we postulate that Trem2^hi^ macrophages promote renal fibrosis by polarizing TECs and fibroblasts towards fibrogenic phenotypes in an SPP1-dependent manner.

## Discussion

The AKI-to-CKD pathological progression represents a coordinated continuum through acute inflammation, resolution and maladaptive fibrotic remodeling. While the inflammatory and fibrotic endpoints have been well characterized, the regulatory mechanisms governing the transition remain incompletely understood. This critical gap prompted us to investigate the immune landscape during this pivotal window. To characterize the dynamic changes of immune cells during AKI-to-CKD transition, we performed scRNA-seq on renal CD45^+^ cells at different timepoints of uIRI models. We identified Trem2^hi^ macrophages as key regulators that bridge inflammation and fibrosis. We provided evidence that Trem2^hi^ macrophages exhibited dual functions: resolving inflammation while promoting fibrosis during AKI-to-CKD transition. By generating Trem2 knockout mice and bone marrow chimeric mice, we found that Trem2 deficiency sustained renal inflammation after uIRI and delayed the progression of fibrosis. This compromised fibrotic programming, while maintaining persistent inflammation under Trem2 deficiency may extend the therapeutic window for interventions against CKD.

Trem2 is a myeloid triggering receptor predominantly expressed on macrophages that senses tissue damage signals and modulates disease progression in various contexts ^33^, including Alzheimer’s Disease, nonalcoholic steatohepatitis and atherosclerosis ^34–36^. Trem2 signaling maintains conserved anti-inflammatory functions across organs, yet its role in fibrosis remains controversial. ScRNA-seq identifies TREM2+CD9+ macrophages as pro-fibrogenic and scar-associated macrophages (SAMs) in human liver ^37^ and lung, expressing *SPP1*, *GPNMB*, *FABP5*, and *CD63* ^38^. These monocyte-derived SAMs localize to fibrotic scars and activate myofibroblasts in response to inflammatory signals. In our study, renal Trem2^hi^ macrophages also highly expressed SAM-related signatures and their deficiency reduced pro-fibrotic factors (Spp1, TGF-β1), supporting its pro-fibrotic role. However, Trem2 function appeared to be context-dependent, as its deficiency exacerbates both inflammation and fibrosis in the unilateral ureteral obstruction (UUO) model-which involves persistent injury stimulation due to continuous obstruction ^39^. Whereas, the self-limited IRI model reveals a more complex role: Trem2^+^ macrophages help resolve acute inflammation but also promote fibrotic remodeling through mediators like Spp1. Furthermore, some studies reported that Trem2 deficiency exacerbates steatohepatitis and liver fibrosis by increasing TGF-β1 secretion ^40^ and impaired collagenolytic activities against CollagenIV ^38,41^. Notably, the composition of the renal fibrotic niche is dominated by CollagenI, which Trem2^hi^ macrophages are unable to degrade. This fundamental difference in injury context and ECM composition underlies the divergent roles of TREM2 signaling across different organs and disease models. In this study, we demonstrated that Trem2 deficiency delayed the initiation of fibrosis and facilitated the sustained inflammation from D3 to D14 after AKI. However, the long-term impact of Trem2 deficiency on inflammation and fibrosis remained determined. It remains the possibility that Trem2^hi^ macrophages exert a time-dependent regulatory role, modulating the pace and intensity of inflammation and fibrosis during the transition from AKI to CKD.

Efferocytosis refers to the engulfment of dying cells by phagocytes, which is crucial for both homeostasis and disease ^42^. In chronic inflammatory diseases, inefficient efferocytosis caused accumulated dead cells and provoked secondary inflammation ^22^. Trem2 has been implicated in microglial phagocytosis of apoptotic neurons and myelin debris ^42^. In renal IRI, tubular injury generates abundant apoptotic or necrotic cells ^44^. We proposed Trem2 mediated macrophage efferocytosis to clear these cells and resolve inflammation. The aberrant accretion of these dying cells may repel the repair process and cause sustained inflammation, as observed in Trem2 deficient uIRI mice. Most mammalian cells can synthesize cholesterol but cannot catabolize. Transporters Abaca1/Abcg1 are required for exporting excessive cholesterol ^23,45^. Trem2 is considered to regulate cholesterol transport and metabolism in microglia and foamy cells ^46,47^. In this study, we found cholesterol accumulation in Trem2KO-BMDMs. LXRs respond to cholesterol overload by upregulating Abaca1/Abcg1 ^48,49^. We found that Trem2 deficiency downregulated LXRs and Abca1/Abcg1, consistent with prior studies ^50^. Trem2^−/−^ foamy cells showed impaired induction of cholesterol efflux genes compared with WT in reported RNA-seq data ^46^. Although some specific oxysterols are known ligands to LXRs, the accumulated oxysterols failed to activate LXRs because of the low LXRs expression in KO-BMDMs. In addition, LXRs activation not only increase cholesterol efflux, but also dampen inflammation by suppressing inflammatory genes and neutrophil migration ^49,50^. We found that the inflammatory status of KO-BMDMs was suppressed by LXR agonist. Our results supported the notion that the inflammatory phenotype of Trem2KO macrophages was caused by low expression of LXR-Abca1/Abcg1 and excess accumulation of cholesterol.

Normal TECs and quiescent fibroblast maintain the structure of healthy kidneys. Upon injury, TECs lose epithelial identity and undergo epithelial–mesenchymal transition ^52^. Meanwhile, myofibroblasts are activated and produce ECM components, leading to renal fibrosis ^53^. Our study revealed that Trem2^hi^ macrophages strongly communicated with pro-fibrotic TECs and myofibroblasts through secreting Spp1. As a soluble factor, Spp1 participates in wound healing by affecting differentiation and activation of myofibroblasts ^54^. However, previous research mostly focused on TECs derived Spp1 ^55,56^. Our data unraveled the involvement of Trem2 signaling in the Spp1 production by macrophages. Trem2^hi^ macrophages derived Spp1 encouraged the pro-fibrotic phenotype of TECs and myofibroblasts activation.

Our study identifies several limitations for future research. First, the upstream signals guiding the differentiation of monocytes into Trem2^hi^ macrophages in injured kidney remain unknown. Second, the mechanism of Trem2-regulated cholesterol metabolism via LXR-ABCA1/ABCG1 needs further investigation. Third, the precise molecular intermediaries in Trem2 downstream signaling require deeper dissection. Fourth, the causal role of macrophage-derived Spp1 in promoting fibrosis requires definitive validation in vivo using cell-specific knockout models. Finally, inducible knockout or timely siRNA approaches during a specific window should clarify whether transient Trem2^hi^ macrophages inhibition can suppress pro-fibrotic responses while preserving their reparative functions, thereby safely halting AKI-to-CKD progression.

In summary, we depict a comprehensive MPCs atlas of AKI-to-CKD transition by scRNA-seq and identify a critical macrophage subset with specific Trem2 expression. These Trem2hi macrophages originated from bone marrow and played a pivotal role in transitioning the kidney from injury and inflammation toward fibrotic remodeling in the uIRI model. Trem2 deficiency resulted in sustained injury and inflammation following AKI, with impaired initiation of fibrotic repair. Mechanistically, Trem2^hi^ macrophages exerted anti-inflammatory effects through efferocytotic clearance of dead TECs and cholesterol metabolism regulated cytokine production. And its pro-fibrotic effect is achieved by activating fibroblasts and TECs trans-differentiation via Spp1. The comprehensive roles that Trem2^hi^ macrophages played in AKI-to-CKD indicated that they orchestrated the transition from acute tubular damage to chronic fibrotic remodeling, deciphering the underlying mechanisms of the inflammatory-fibrosis axis. (Fig 8). These findings highlight the functional complexity of macrophages during the transition from inflammation to fibrosis. Targeting Trem2 may delay fibrosis onset and extend the therapeutic window for promoting inflammation resolution and tubular repair, if appropriately timed strategies are employed. This work provides a translational basis for developing integrated therapeutic approaches that concurrently modulate immune responses and enhance tissue repair to halt the progression from AKI to CKD.

**Figure 8.**
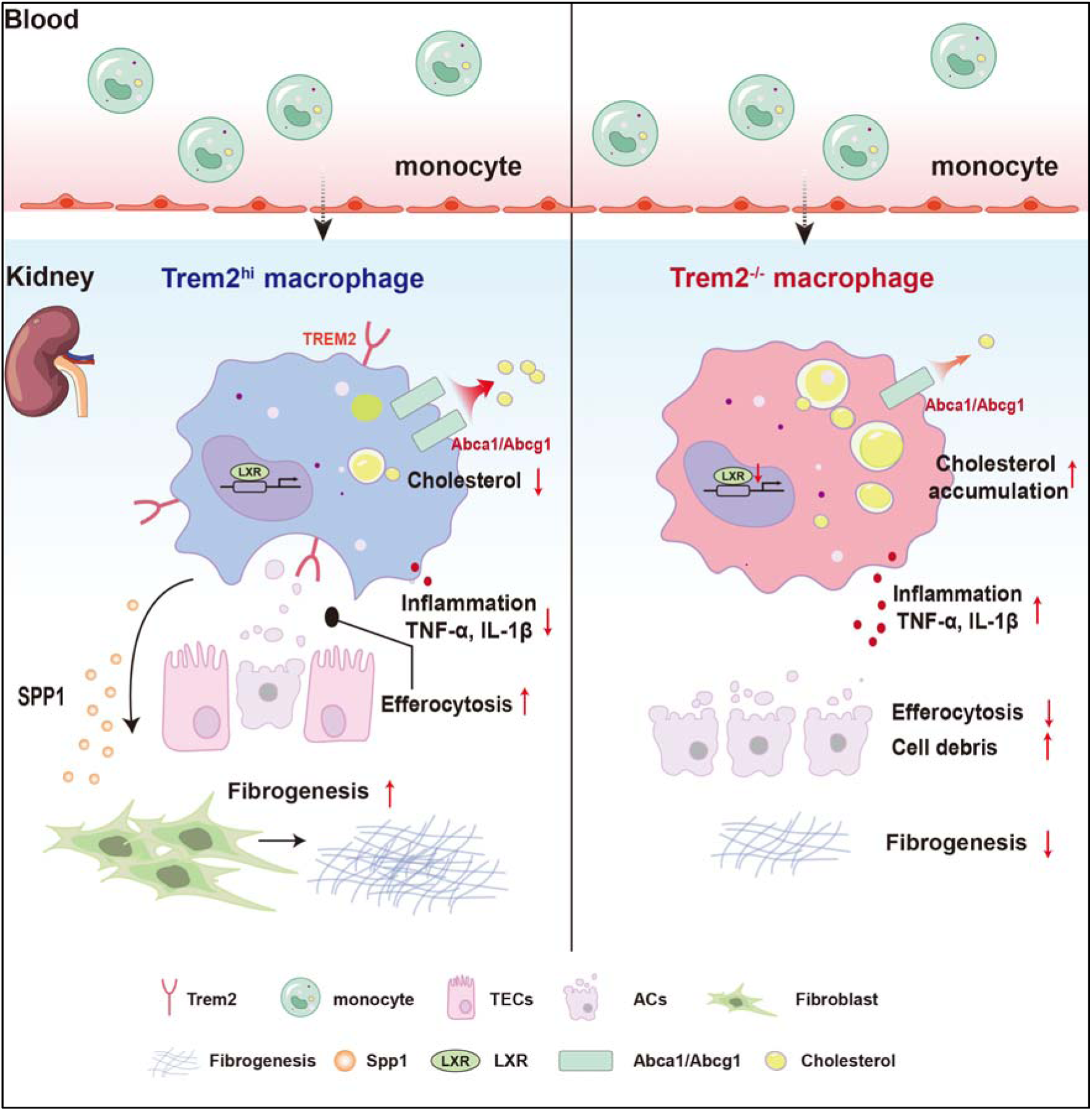
Schematic diagram showing the function of Trem2^hi^ macrophages in AKI-to-CKD transition. Time-course analysis revealed that acute ischemic-reperfusion injury triggered the recruitment of bone marrow-derived Trem2^hi^ macrophages to the injured kidney. These macrophages contributed to the resolution of inflammation through efferocytic clearance of apoptotic tubular epithelial cells (TECs) and suppression of pro-inflammatory cytokine release via Lxr-Abcg1/Abca1-dependent cholesterol efflux. Additionally, Trem2^hi^ macrophages produced Spp1 to promote TEC trans-differentiation and myofibroblast activation, facilitating extracellular matrix production and kidney remodeling. Genetic deletion of Trem2 resulted in persistent inflammation and impaired initiation of fibrotic remodeling, due to the accumulation of cellular debris and sustained cytokine production following kidney injury.

## Data Statement

The scRNA-seq data supporting the findings of this study are publicly available in Gene Expression Omnibus at https://www.ncbi.nlm.nih.gov/geo/, reference number GSE240885.

## Authors’ contributions

Yan Tong, Fangxin Mu and Chao Wang, literature research, conception and design, conducting experiments, validation, acquiring data, data interpretation and writing manuscript. Xue Wang and Tian Sang, conducting experiments, investigation and data visualization. Yulan Chen and Jian Zhang, conducting experiments and acquiring data. Xu Wang, molecular experiments assistance. Ran Liu, animal experiments assistance. Shaoyuan Cui and Bo Fu, cellular experiments assistance. Jiaona Liu, pathological experiments assistance. Li Zhang, Xueyuan Bai, Qinggang Li, design guidance and processing data. Quan Hong, Xuefeng Sun, Guangyan Cai, conception and design guidance, supervision and revising manuscript. Qing Ouyang and Xiangmei Chen, funding acquisition, conception and design, data interpretation, supervision and revising manuscript.

## Acknowledgments

This work was supported by the National Natural Science Foundation of China (NO. 82570797, 82000657, 82030025 and 32141005). We thank Beijing EASYRESEARCH Co., Ltd for the support of bioinformatic analyses.

