## Supplementary materials for "Trem2^hi^ macrophages bridge inflammation resolution and fibrosis initiation after ischemia-reperfusion injury in the kidney"

*Yan Tong et al.*

**This PDF file includes:**

Supplementary Text

Figs. S1 to S7

Tables S3

**Methods**

**Tissue dissociation and preparation of single-cell suspension**

Murine kidneys were freshly obtained at the indicated timepoints from anesthetized mice and cut into small pieces on ice, and then digested in RMPI 1640 medium containing 1 mg/mL Collagenase I, 1mg/mL Collagenase IV and 100 μg/mL DNase I (Sigma-Aldrich) at 37℃ for 30 min using an MACSmix™ Tube Rotator (Miltenyi Biotec). The digested tissue was then passed through a 70 μm and 40 μm cell strainers (Falcon, BD Biosciences) to remove debris. After washing in pre-chilled PBS, the cell pellets were incubated with 1× red blood cell (RBC) lysis buffer (Miltenyi Biotec) on ice for 5 min. Then the cell pellets were washed twice to obtain single-cell suspension.

**scRNA-Seq data processing and analysis**

Demultiplex cellular barcodes and raw sequencing data were processed using Cell Ranger software pipeline (version 5.0.0) provided by 10× Genomics. STAR aligner was used to map reads to genomes and transcriptomes across samples and produce a matrix of gene counts versus cells. The R package Seurat (Version 4.3.1) was employed to perform Unique Molecular Identifier (UMI) counts of single cells analysis, normalization, dimension reduction and many other functions below. For quality control criteria, cells with UMI more than 1000, gene counts between 200 to 1000 and mitochondrial gene percentages less than 20% were kept and potential cell doublets and multiple captures were discarded. The filtered count matrix was then normalized and scaled by using NormalizeData function in Seurat to generate normalized gene counts with method “Log Normalize”. The downstream analyses were conducted based on these cells.

Top 2000 variable genes across filtered single cells were screened using the FindVariableGenes and their normalized expression were scaled with percentage of mitochondrial RNA and gene numbers regressed out. To mitigate batch effects across individuals, integration anchors were computed between datasets using the canonical correlation analysis (CCA) algorithm embedded in Seurat. These anchors were input into the IntegrateData function, enabling harmonized downstream analysis ^58^. For visualization, the dimensionality was further reduced using Uniform Manifold Approximation and Projection (UMAP) or T-Distributed Stochastic Neighbor Embedding (t-SNE). The FindClusters function based on the gene expression profiles of the cell was used to perform graph-based clustering. The whole dataset and MPC subset were clustered at resolutions of 1.0 and 0.3, respectively. We used the identified cell type specific expressed signatures calculated by the FindAllMarker function, combined with canonical marker genes in previously published literature to annotate each cell type. For the visualization of gene expression, scatter plots, dot plots, violin plots and volcano plots were implemented through the FeaturePlot and VlnPlot functions. RNA velocity analysis was performed by using velocyto CLI version 0.17. Individual sample BAM files along with a mouse gene annotation file (GTF) for mm10-3.0.0 were input to the ‘run 10×’ command. To estimate and visualize RNA velocity vector fields in Seurat integrated UMAP space, splicing kinetics model was built by scVelo version 0.2.4 ^59^. P value＜0.01 and∣log_2_Fold Change∣＞0.9 were set as thresholds for significant DEG expression. KEGG enrichment analysis of DEGs was performed by R software based on the hypergeometric distribution. ECM gene sets defined by the Matrisome Project were applied for ECM scoring ^19^.

**Flow cytometry analysis**

Single-cell suspensions of kidney were prepared as described above. For cell surface marker staining, single cell suspensions were incubated with antibodies for 30 min at 4℃. For intracellular marker staining, single cell suspensions were fixed and permeabilized using the Foxp3/Transcription Factor Staining Buffer Set (eBioscience), followed by antibodies staining for 30 min at 4℃. The samples were washed and acquired on BD LSRFortessaTM cytometer. The data were analyzed by FlowJo V.10.0.7 software. Most of the antibodies and isotype controls used in this study were purchased from BioLegend: Zombie NIR™ Fixable Viability Kit, TruStain FcX™ CD16/32, CD45-PE, F4/80-BV421, CD11b-PerCp/Cy5.5, CD206-APC, CD45.2-PE, CD45.1-BV785, CD4-FITC and CD19-BV421, CD279-APC, CD8a-BV605, T-bet-PE/Cy7, PD-1-APC. Other antibodies were Trem2-FITC, Gpnmb- eFluor™ 660, Foxp3-PE and CD45-PerCp/Cy5.5.

**Isolation and Culture of Bone Marrow Derived Macrophage (BMDM)**

Bone marrow (BM) cells were isolated from the femurs and tibias of 6-8 weeks old male C57BL/6J mice. The cell suspension was then passed through a 70 μm filter, followed by RBC lysis. BM derived macrophages (BMDMs) were induced and cultured in 1640 RMPI medium containing 10% FBS (Fetal Bovine Serum), 1% P/S (Penicillin-Streptomycin) (Gibco) and 20 ng/ml M-CSF (Macrophage Colony Stimulating Factor) (Peprotech) at 37℃ and 5% CO2 levels for 3 days before changing the medium for the first time. Differentiated or stimulated BMDMs were harvested for further experiments at day 7.

**Induction of apoptosis of tubular cells and efferocytosis assay**

For induction of apoptosis, Mouse kidney tubular cells (TCMK-1 cell line) were treated with 10 μM rotenone for 24 h. The apoptosis rate was determined by staining AV/PI (Annexin V / Propidium Iodide) (BioLegend, #640914). This method routinely yielded more than 85% Annexin V^+^ cells. For efferocytosis experiment, ACs were stained with PHrodo dye (ThermoFisher, #P36011) for 30 min at room temperature. BMDMs from WT and Trem2KO mice were cultured in plates and incubated with PHrodo-labeled ACs. After 1 h, unbound ACs were vigorously washed off, and BMDMs were stained with F4/80. The engulfment rate was determined as the percentage of PHrodo-positive in the F4/80^+^ cell population by flow cytometric analysis.

**Cholesterol detection**

Lipids in cells were stained with F4/80 and BODIPY (Boron-Dipyrromethene) Lipid Probe (ThermoFisher, #D3921) for 30 min and detected by flow cytometry. For cholesterol staining, cells were stained with Filipin III and observed by an Olympus confocal microscope (Olympus).

Cholesterol and its metabolites were detected by using electrospray ionization (ESI) mode on the Exion UPLC QTRAP 6500 PLUS Sciex) LC/MS (LipidALL Technologies). Oxysterols and sterols were quantitated by referencing the spiked internal standards and normalized to total protein in BMDM cells.

**Cellular fibrosis assays**

Conditioned media (CM) from ACs-phagocytosing Trem2 WT/KO BMDMs were generated according to the following protocol. Briefly, BMDMs from WT and Trem2KO mice were cultured in plates and incubated with ACs for 24h. Subsequently, the media were replaced with fresh media and incubated for another 48h. The supernatants were then collected, centrifuged to remove cellular debris for further use. These media were used as stimulation for primary tubular epithelial cells or 3T3 fibroblasts. And the concentrations of Spp1 in these WT-CM and KO-CM were measured by ELISA. For blockade experiment, the Spp1 neutralization antibody (Sigma-Aldrich, #O7635) was used at a concentration of 2 μg/mL.

**Enzyme-linked immunosorbent assays (ELISA)**

The level of IL-1β, TNF-α and Spp1 in cell culture supernatants were quantified using commercial mouse ELISA kits (RayBiotech, # ELM-IL1b, # ELM-TNFa; Bio-Techne, #MOST00) according to the manufacturer’s instructions. In brief, standards and samples were added to the antibody-precoated microplate and incubated for 2.5 h at room temperature. The plate was washed four times, followed by incubation with a biotinylated detection antibody for 1 h. After repeated washing, prepared streptavidin solution was added and incubated for 45 min. Following a final wash, TMB substrate was applied and incubated for 30 min under protection from light. The reaction was terminated with stop solution, and the absorbance was measured using a microplate reader set to 450 nm. Sample concentrations were determined based on a standard curve generated with a four-parameter logistic model.

**Renal function measurement**

To evaluate the renal function of AKI-to-CKD model, the levels of serum creatinine and blood urea nitrogen at the indicated timepoints were detected using the QuantiChrom Creatinine and Urea Assay kits (DICT-500, DIUR-500, BioAssay Systems), respectively according to the manufactural instructions.

**Renal tissue histopathological analysis**

Murine kidney samples were fixed in 10% formalin and then paraffin-embedded sections (3 μm) were stained with periodic acid-Schiff (PAS) and Masson trichrome staining. The degree of tubular injury and interstitial fibrosis were assessed and scored by two experienced pathologists blinded to the experimental groups using an Olympus BX53 microscope equipped with a digital camera (Olympus).

Ten random microscopic fields of each section were selected for analysis with a 20x objective lens. For PAS staining, we evaluated the tubular injury and necrosis using a method referring to ^60^ as follows: 0 = no damage, 1 = brush border loss, tubular cell swelling with less than 1/3 tubular cells showing nuclear loss, 2 = brush border loss, tubular cell swelling with less than 1/3—2/3 tubular cells showing nuclear loss, 3 = more than 2/3 tubular cells showing nuclear loss. Fibrosis progression was assessed through Masson's trichrome staining by calculating the percentage of collagen deposition relative to total field area. Histopathological scoring was performed according to established criteria ^61^ as follows: Grade 0 - no detectable fibrosis; Grade 1 - fibrosis <25% of field area; Grade 2 - fibrosis accounting for 25-50% of microscopic field; Grade 3 - fibrosis accounting for 50-75% of microscopic field; Grade 4 - fibrosis accounting for >75% of microscopic field.

Immunofluorescent staining. After fixation with 4% paraformaldehyde, murine kidney tissue cryosections (4 μm) were permeabilized with 1% Triton X-100 (Sigma-Aldrich) for 10 min and then blocked with 1% bovine serum albumin (BSA, Sigma-Aldrich) for 1 h at room temperature. Sections were incubated with primary antibodies at 4℃ overnight, followed by appropriate secondary antibodies incubation for 1 h at room temperature. Cell nuclei were stained with DAPI when mounting the slides. All fluorescence images were acquired using a confocal laser microscope (FV10-ASW, Olympus). All the images were analyzed using ImageJ software and calculated from 10 random and individual high-power fields. The primary antibodies in this experiment were as follows: α-SMA (Abcam, #ab7817), Fibronectin1 (Proteintech, #15613-1-AP), LAMP (Santa-Cruz, #sc-19992).

TUNEL staining was performed in kidney sections using TUNEL assay kit (Beyotime, #C1090) according to the manufactural instructions.

**Quantitative Real-Time PCR (qRT-PCR)**

Total RNA from kidney tissue and cells were isolated using the TRIzol method. RNA concentration and purity were detected by Nanodrop. Total RNA was reverse-transcribed into cDNA using a PrimeScript reverse transcription reagent (Takara, #6215A). An SYBR Green PCR amplification was performed by real-time PCR system (ABI QuantStudio5 Q5). The relative mRNA expression levels were calculated and normalized to house-keeping gene β-actin. Primer sequences are presented in Table S3.

**Western Blotting analysis**

Kidney tissue and cells were lysed in lysis buffer (Beyotime, #P0013) containing protease and phosphatase inhibitors (Solarbio, #P6731). Protein samples were electrophoresed and separated on SDS-PAGE, and then transferred onto nitrocellulose filter membrane using a Turbo Transfer System (Bio-Rad). The membranes were blocked with 5% skim milk for 2 h at room temperature. Then the membranes were incubated with primary antibodies at 4℃ overnight. After washed with Tris-buffered saline-Tween 20, the membranes were incubated with HRP-conjugated secondary antibodies for 2 h at room temperature. Protein bands were visualized using an enhanced chemiluminescence (ECL) system (Bio-Rad). The primary antibodies in this study were as follows: Trem2 (CST, #76765), CollagenI (Proteintech, #14695-1-AP), Vimentin (Abcam, #ab92547), Fibronectin1 (Proteintech, #15613-1-AP), Rab5a (Santa-Cruz, #sc-515401), Rab7 (Santa-Cruz, #sc-271608), GAPDH (Proteintech, #10494-1-AP), β-actin (Proteintech, #20536-1-AP). The secondary antibodies were purchased from Beyotime.

### Supplementary Figures

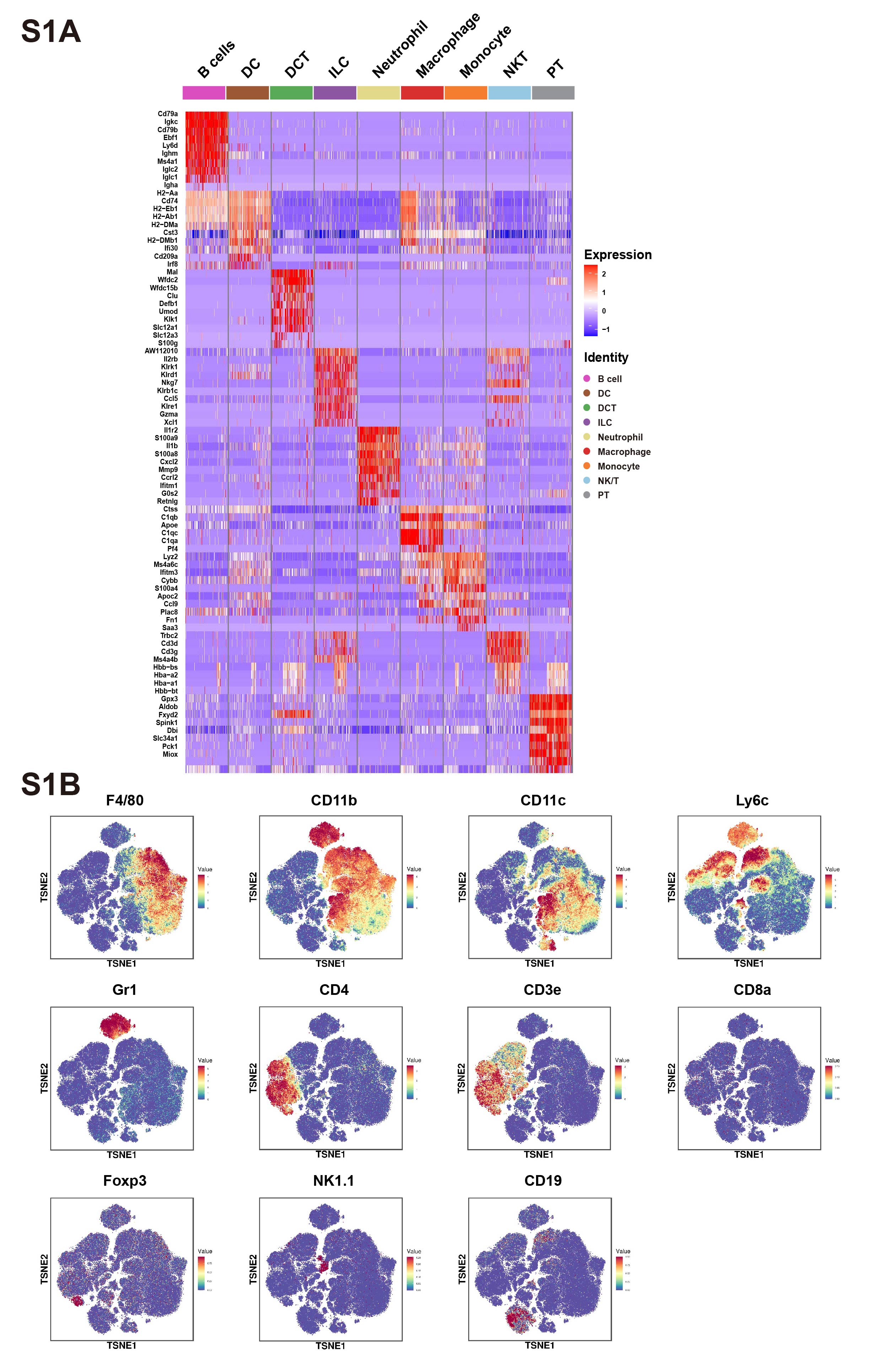

**Fig S1 The feature of cell types and cell counts by single-cell RNA sequencing and CyTOF.**

(A) Heatmap of the top 10 genes of each cell type by single-cell RNA sequencing.
(B) Feature plots of immune cells by CyTOF.

PT, proximal tubule; DCT, distal convoluted tubule; DC, dendritic cell; ILC, innate lymphoid cell. CyTOF, cytometry by time of flight.

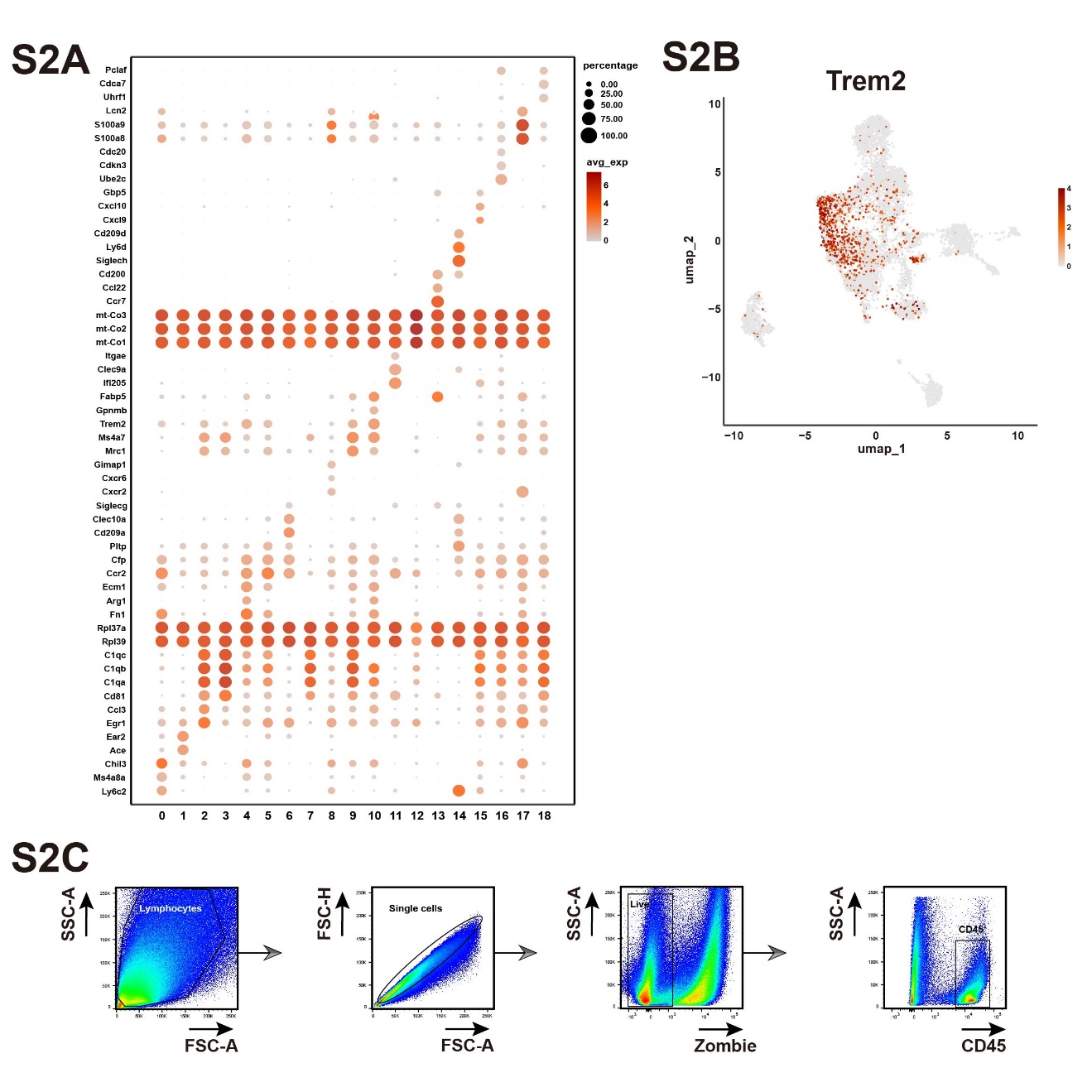

**Fig S2 The marker genes of MPC cluster and the gating strategy for macrophage.**

(A) Dot plot of the marker gene expression of cells in each MPC cluster.
(B) The gene expression of Trem2 in MPCs.
(C) The flow cytometry gating strategy for kidney macrophages.

MPC, mononuclear phagocytic cell.

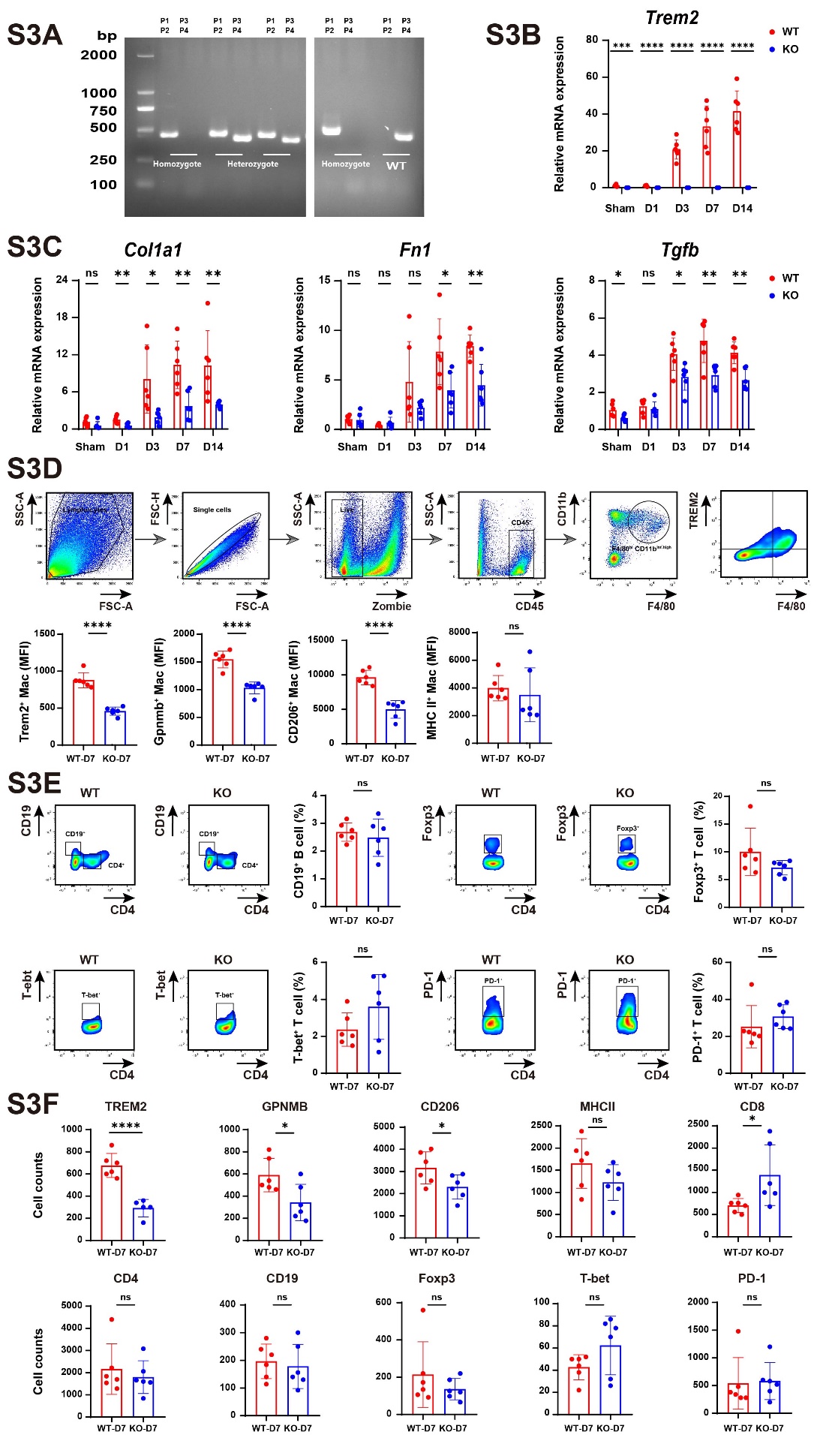

**Fig S3 The mRNA expression levels of fibrotic markers and percentages of other immune cells in WT and KO mice after uIRI.**

(A) Verification of Trem2 gene deletion by PCR using primer pairs P1/P2 and P3/P4.
(B) Verification of Trem2 gene deletion by RT-qPCR.
(C) The mRNA expression levels of *Col1a1*, *Fn1* and *Tgfb* detected by RT-qPCR.
(D) The gating strategy and the quantitative data of mean fluorescence intensity (MFI) of macrophages by flow cytometry.
(E) The percentages of other immune cells in WT and KO mice at D7 were analyzed by flow cytometry.
(F) The absolute number of specific subsets within per 200,000 total cells in WT and KO mice at D7 were analyzed by flow cytometry.

Wild type: PCR with P1 and P2 cannot amplify any fragment; PCR with P3 and P4 can obtain a 415bp fragment.

Heterozygote: PCR with P1 and P2 yields a 449bp fragment; PCR with P3 and P4 can obtain a 415bp fragment.

Homozygote: PCR with P1 and P2 yields a single 449bp fragment; PCR with P3 and P4 cannot obtain any bands.

Data are presented as the mean±SD, n=6, **P*＜0.05, ***P*＜0.01, ****P*＜0.001, *****P*＜0.0001, ns, no significance.

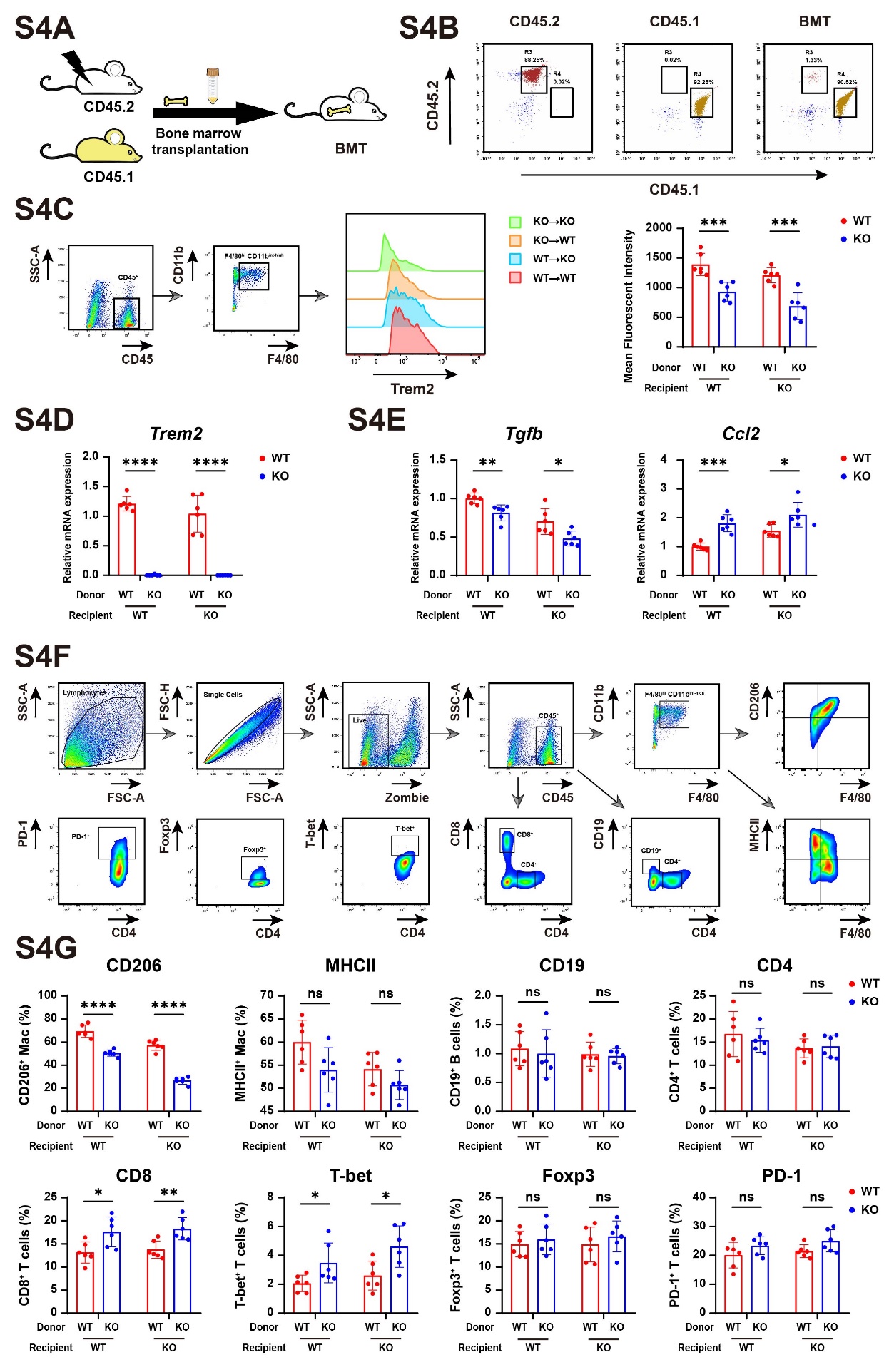

**Fig S4 The construction of BMT model and percentages of other immune cells in BMT mice after uIRI.**

(A) Experimental protocol of BMT to elucidate the origin of Trem2^hi^ macrophages.
(B) The CD45.1 or CD45.2 expression of immune cells in peripheral blood were analyzed by flow cytometry to demonstrate the successful BMT model.
(C) The mean fluorescent intensity of Trem2 were analyzed by flow cytometry.
(D) The mRNA expression levels of *Trem2* in BMT mice detected by RT-qPCR
(E) The mRNA expression levels of *Tgfb* and *Ccl2* in BMT mice detected by RT-qPCR.
(F) The gating of other immune cells in BMT mice at D7 were analyzed by flow cytometry.
(G) The percentages of other immune cells in BMT mice at D7 were analyzed by flow cytometry.

Data are presented as the mean±SD, n=6, **P*＜0.05, ***P*＜0.01, ****P*＜0.001, *****P*＜0.0001, ns, no significance.

BMT, bone marrow transplantation.

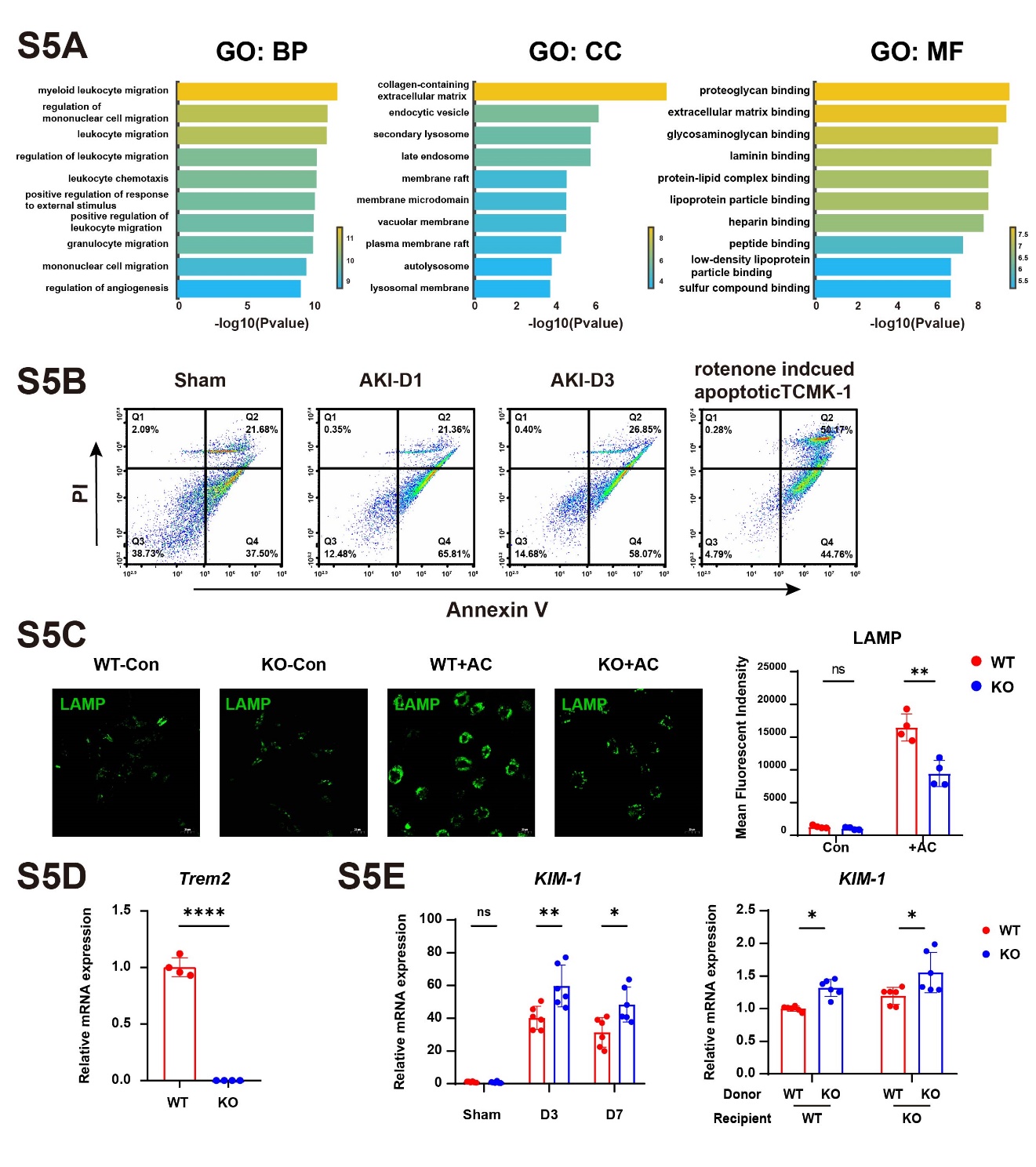

**Fig S5 GO analysis of Trem2^hi^ macrophages and apoptotic cells in vivo and in vitro.**

(A) GO analysis based on the differentially expressed genes of Trem2^hi^ macrophages.
(B) The proportions of apoptotic cells in sham kidney and injured kidneys at D1 and D3 as well as rotenone induced apoptotic TCMK-1 cells.
(C) The LAMP staining of BMDMs from WT and KO mice in the absence or presence of ACs.
(D) Relative mRNA expression levels of *Trem2* in BMDM from WT or KO mice was measured by RT-qPCR.
(E) Relative mRNA expression levels of *KIM-1* in kidney from WT or KO mice was measured by RT-qPCR.

Data are presented as the mean±SD, n=6, **P*＜0.05, ***P*＜0.01, ****P*＜0.001, *****P*＜0.0001, ns, no significance.

TCMK-1 mouse kidney tubular cell line. LAMP, lysosome-associated membrane proteins. BMDM, bone marrow derived macrophages.

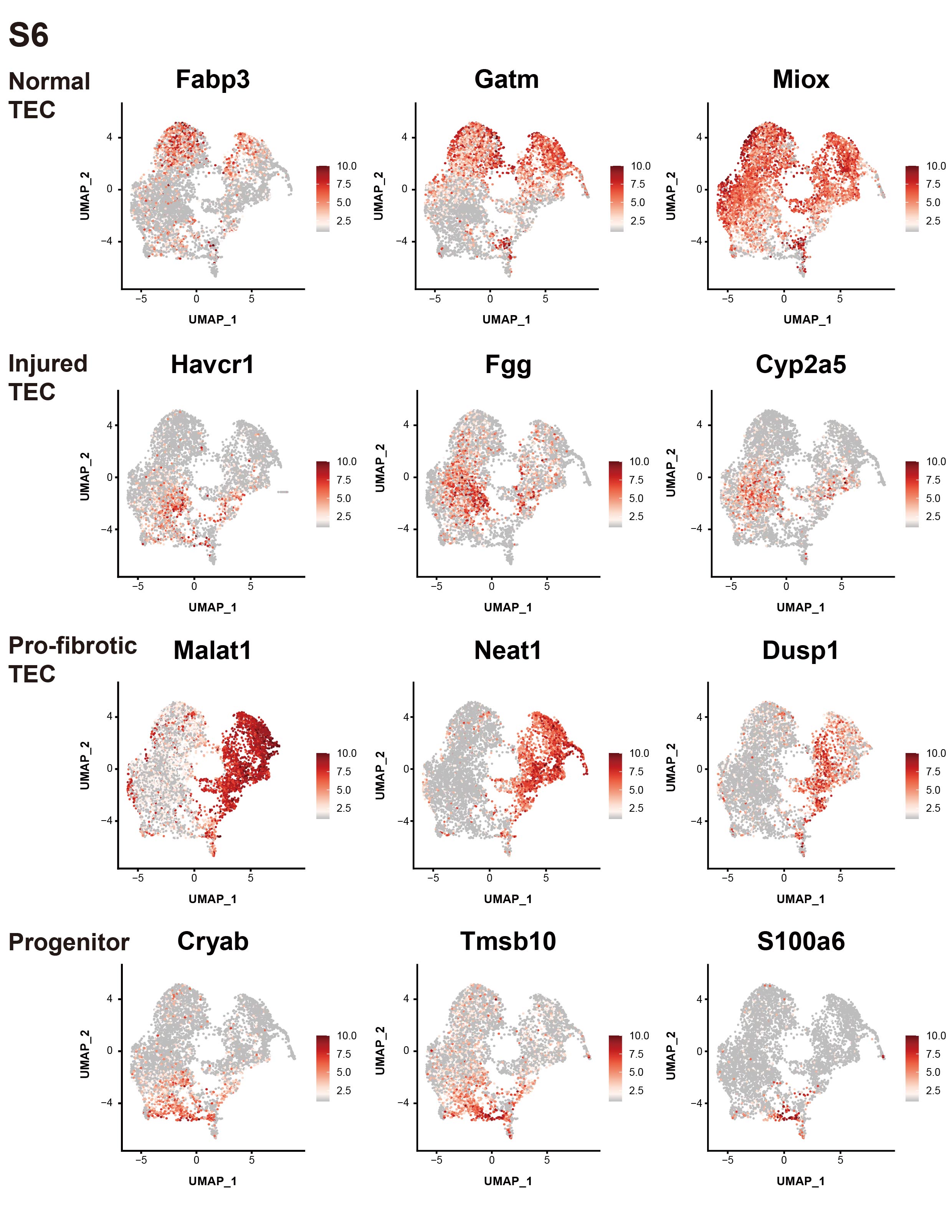

**Fig S6 Umap plot of the marker genes in TEC sub-clusters. TEC, tubule epithelial cell.**

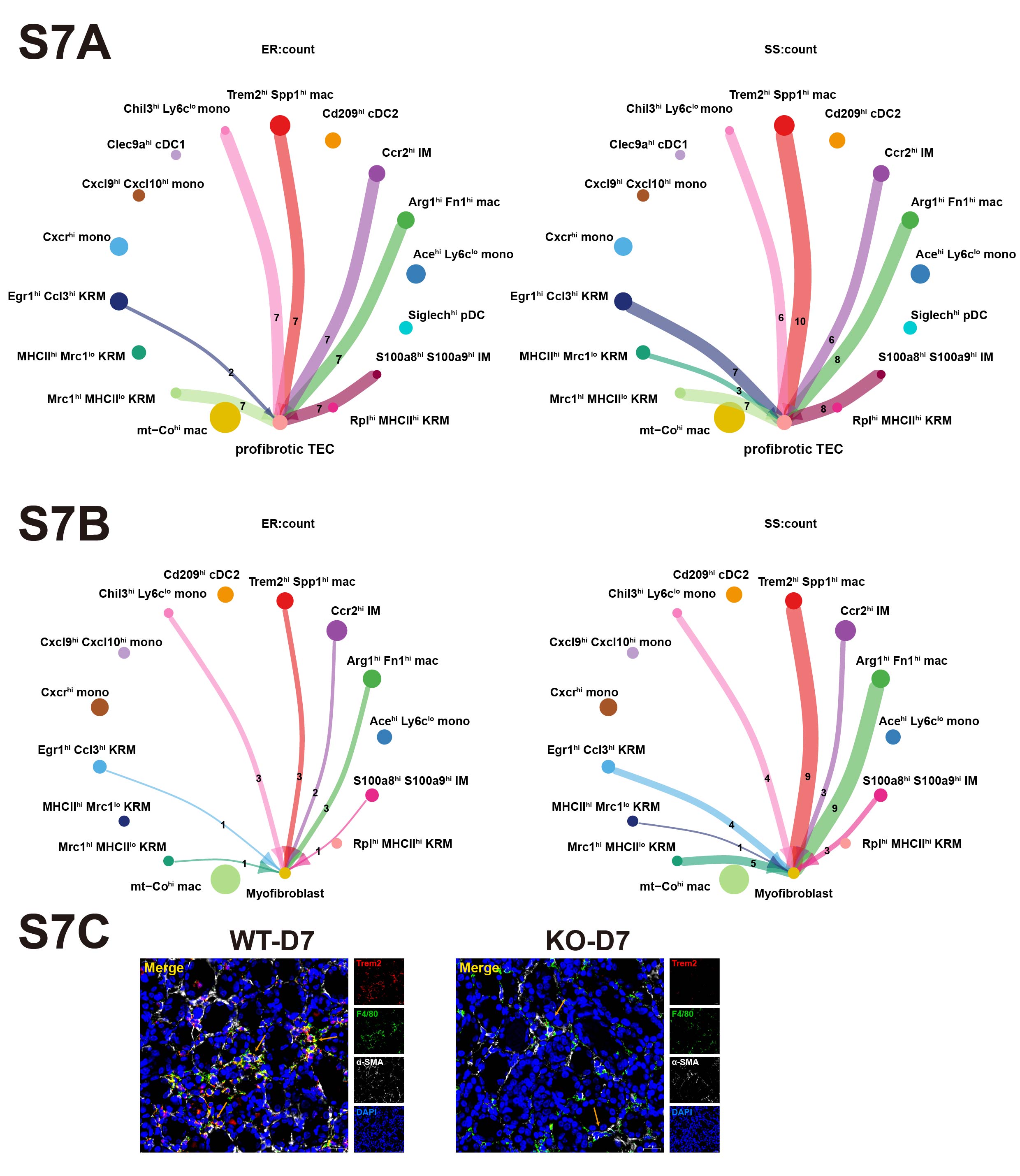

**Fig S7 Interactions between Trem2+ macrophages and tubule cells or fibroblasts.**

(A) The circle plots showing the numbers of interactions between all macrophage subtypes and pro-fibrotic TECs.
(B) The circle plots showing the numbers of interactions between all macrophage subtypes and myofibroblasts.
(C) Representative immunofluorescence images of Trem2^+^ macrophages (F4/80^+^ Trem2^+^) that co-localize with fibrotic tubular cells or myofibroblasts (α-SMA^+^) in WT-D7 and KO-D7 kidney.

**Table S3**

List of Primers for RT-qPCR

| Gene | Forward Primer | Reverse Primer |
| --- | --- | --- |
| *Actb* | GTGACGTTGACATCCGTAAAGA | GCCGGACTCATCGTACTCC |
| *Trem2* | ATCGCAGATGACACCCTTGC | CTTGGGCACCCTCGAAACT |
| *Spp1* | GAGGAAACCAGCCAAGGACT | AAGCTTCTCCTCTGAGCTGC |
| *Gpnmb* | GGCTACTTCAGAGCCACCATCA | CTTTGCAGGTCACAGTGAAGTCC |
| *Ctsd* | TAAGACCACGGAGCCAGTGTCA | CCACAGGTTAGAGGAGCCAGTA |
| *Col1a1* | GCTCCTCTTAGGGGCCACT | ATTGGGGACCCTTAGGCCAT |
| *Fn1* | TCCCAGAGAAGTGGTCCCTC | TGGGGAAGCTCATCTGTCTT |
| *Tgfb* | CTGATACGCCTGAGTGGCTG | TTTGGGGCTGATCCCGTTG |
| *Il-1b* | GAAATGCCACCTTTTGACAGTG | TGGATGCTCTCATCAGGACAG |
| *Ccl2* | TTAAAAACCTGGATCGGAACCAA | GCATTAGCTTCAGATTTACGGGT |
| *Tnfa* | CAGGAGGGAGAACAGAAACTCCA | CCTGGTTGGCTGCTTGCTT |
| *Hmgcr* | AGCTTGCCCGAATTGTATGTG | TCTGTTGTGAACCATGTGACTTC |
| *Hmgcs1* | AACTGGTGCAGAAATCTCTAGC | GGTTGAATAGCTCAGAACTAGCC |
| *Srebp2* | GTGCGCTCTCGTTTTACTGAAGT | GTATAGAAGACGGCCTTCACCAA |
| *Acat1* | CAGGAAGTAAGATGCCTGGAAC | TTCACCCCCTTGGATGACATT |
| *Lxra* | CTCAATGCCTGATGTTTCTCCT | TCCAACCCTATCCCTAAAGCAA |
| *Lxrb* | ATGTCTTCCCCCACAAGTTCT | GACCACGATGTAGGCAGAGC |
| *Abca1* | GGTTTGGAGATGGTTATACAATAGTTGT | CCCGGAAACGCAAGTCC |
| *Abcg1* | CTTTCCTACTCTGTACCCGAGG | CGGGGCATTCCATTGATAAGG |
